# Computational modeling of TGF-β2:TβRI:TβRII receptor complex assembly as mediated by the TGF-β co-receptor betaglycan

**DOI:** 10.1101/2022.05.09.490905

**Authors:** Michelle Ingle, Asakiran Madamanchi, Andrew P. Hinck, David Umulis

**Affiliations:** Department of Agricultural and Biological Engineering, Purdue University, West Lafayette, IN 16802; Department of Structural Biology, University of Pittsburgh School of Medicine, Pittsburgh, PA 15260

**Author notes:** Current Address: School of Information, University of Michigan, Ann Arbor, MI, 48109. Corresponding Author: Prof. David Umulis, Department of Agricultural and Biological Engineering, Weldon School of Biomedical Engineering, Purdue University West Lafayette, IN 47907, U.S.A.

**Keywords:** TGF-β, betaglycan, TβRIII, TβRII, TβRI, computational modeling, receptor complex assembly

## Abstract

Transforming growth factor-β1, -β2, and -β3 (TGF-β1, -β2, and -β3) are secreted signaling ligands that play essential roles in tissue development, tissue maintenance, immune response, and wound healing. TGF-β homodimers signal by assembling a heterotetrameric complex comprised of two type I receptor (TβRI):type II receptor (TβRII) pairs. TGF-β1 and TGF-β3 signal with high potency due to their high affinity for TβRII, which engenders high affinity binding of TβRI through a composite TGF-β:TβRII binding interface. However, TGF-β2 binds TβRII 200-500 more weakly than TβRII and signals with lower potency compared to TGF-β1 and -β3. Remarkably, potency of TGF-β2 is increased to that of TGF-β1 and -β3 in the presence of an additional membrane-bound co-receptor, known as betaglycan (BG), even though betaglycan does not directly participate in the signaling mechanism and is displaced as the signaling receptors, TβRI and TβRII, bind. To determine the role of betaglycan in the potentiation of TGF-β2 signaling, we developed deterministic computational models with different modes of betaglycan binding and varying cooperativity between receptor subtypes. The models, which were developed using published kinetic rate constants for known quantities and optimization to determine unknown quantities, identified conditions for selective enhancement of TGF-β2 signaling. The models provide support for additional receptor binding cooperativity that has been hypothesized, but not evaluated in the literature. The models further showed that betaglycan binding to TGF-β2 ligand through two domains provides an effective mechanism for transfer to the signaling receptors that has been tuned to efficiently promote assembly of the TGF-β2(TβRII)_2_(TβRI)_2_ signaling complex.

## Introduction

Transforming growth factor-β (TGF-β) signaling is mediated by three secreted ligands, TGF-β1, -β2, and -β3. The TGF-β isoforms share 70 – 80% sequence identity and adopt similar homodimeric structures, consisting of two identical cystine-knotted protomers tethered together by a single inter-chain disulfide bond (1,2). TGF-β signal transduction is activated when the TGF-β type I and type II receptors, TβRI and TβRII, are brought together by ligand binding to initiate a trans-phosphorylation reaction that results in phosphorylation of cytoplasmic Smad2 and Smad3, the canonical effectors of the pathway. Phospho-Smad2 and -Smad3, after forming a heterotrimeric complex with Smad4 (3), translocate to the nucleus where they regulate gene expression in partner with other co-activators and co-repressors (4).

TGF-β1 and -β3 orchestrate assembly of their signaling complexes in a step-wise manner, first by binding TβRII with high affinity to form a stable binary complex with TβRII (5). The TGF-β:TβRII binary complex recruits TβRI, to form a highly stable ternary complex, consisting of one ligand homodimer bound to two molecules of TβRI and two molecules of TβRII (6,7).

Though subsequent studies have shown that a TGF-β(TβRI:TβRII)_1_ heterodimer is capable of signaling, the most efficient signaling comes from a TGF-β(TβRI:TβRII)_2_ heterotetramer (8).

These cell-based observations are supported by binding and structural studies with purified TGF-βs and receptor ectodomains, which demonstrate that TβRII binds symmetrically to the fingertips of the extended TGF-β monomers with high affinity and TβRI is recruited by binding a composite interface formed by both TGF-β and TβRII (9–13). The potentiation of TβRI binding by TβRII, which is mediated by direct contact between the TβRII N-terminal tail and TβRI (12), is difficult to quantitate owing to the very weak binding of TβRI by TGF-β but has been estimated to be as high as 10^3^-fold (13).

TGF-β2 binds TβRII about 200 times more weakly than TGF-β1 and -β3 (9,14–16) and in cells lacking the TGF-β type III receptor, also known as betaglycan (BG), its potency is reduced accordingly (9,15,17). BG is comprised of a large (ca. 750 aa) ectodomain, a single-spanning transmembrane domain, and a short (42 aa) non-catalytic cytoplasmic tail and is considered a co-receptor as it binds TGF-βs and potentiates receptor complex assembly and signaling, but does not directly participate in the phosphorylation-mediated signaling cascade (17,18). BG binds all three TGF-β isoforms with high affinity (K_D_ 5 – 20 nM) (19), though its potentiation of receptor complex assembly and signaling, which restores the potency of TGF-β2 to that of TGF-β1 and -β3 (17,20), is greatest for TGF-β2. BG knockout mice are embryonic lethal (21) and share many of the phenotypic characteristics of the TGF-β2 knockout, including pronounced cardiac defects, confirming BG’s importance for TGF-β2 signaling *in vivo* (22).

The extracellular domain of BG is comprised of two subdomains, the N-terminal membrane-distal orphan domain and the membrane-proximal zona pellucida, or ZP domain (19,20). The structure of the BG orphan domain, BG_O_, consists of two tandem β-sandwich domains connected together by a semi-rigid two stranded β-sheet (23). The BG zona pellucida domain, BG_ZP_, is likely comprised of tandem immunoglobulin-like domains, designated BG_ZP-N_ and BG_ZP-C_, based on available structures of BG_ZP-C_ from rat (24) and mouse (25) and homology to the zona pellucida domain of endoglin (26), a homologous co-receptor of the TGF-β family that binds and promotes signaling of BMP-9 and -10.

BG has been suggested, based on crosslinking studies with radiolabeled TGF-β2, to potentiate assembly of the assembly of the signaling complex by a handoff mechanism, in which BG initially captures TGF-β2 on the cell surface, and after promoting binding of TβRII to form a stable ternary complex, is displaced as TβRI binds (17). More recent studies with purified full-length rat BG ectodomain (rBG_O-ZP_), and its subdomains, have shown that BG binds TGF-β2 dimers with 1:1 stoichiometry, with the orphan and ZP-C domain (rBG_O_ and rBG_ZP-C_, respectively) both directly binding the growth factor (19,27). BG-binding blocks one of the TβRII binding sites through its BG_ZP-C_ domain, but leaves the other accessible, and has been proposed to promote binding of TβRII both by membrane-localization effects, and to a lesser extent allostery (27). Binding of full-length betaglycan extracellular domain was shown to interfere with binding of TβRI in the presence of TβRII (27), suggesting that the recruitment of TβRI is responsible for displacing BG as the signaling complex is formed.

Previous studies, while providing a qualitative understanding of receptor complex assembly and signaling, leave unanswered several questions regarding the assembly mechanism. Chief among these is the relative importance of weak interactions, and alternative assembly pathways, that potentially become significant after the initial binding event that captures the ligand on the surface of the membrane. Other questions relate to the apparent paradox of how BG potentiates assembly of the signaling complex by binding TGF-βs with near nanomolar affinity, but is ultimately displaced as the signaling receptors bind. In order to address these questions we used deterministic computational modeling to assess complex assembly and signal generation based on published kinetic rate constants for as many of the steps as possible. In cases, where information was lacking, the values were estimated based on simulations over a range of plausible values. Results suggest that for BG-independent assembly, the interaction between the TβRI with TβRII ectodomains is critical for driving assembly, especially TGF-β2. In addition, the results suggest that it is important to consider alternative pathways for receptor addition, especially for TGF-β2 which binds TβRI and TβRII with comparable weak affinity. Results for BG-dependent assembly reiterate the points noted above, but also suggest that BG’s high affinity, achieved through bivalent binding, is critical for efficient handoff to the signaling receptors.

## Results

### Receptor-receptor interactions and the SRR, TRR, and NRR models for assembly

The importance of the interaction between TβRI and TβRII for the initial recruitment of TβRI into the TGF-β signaling complex has been heavily studied (8,11–13,28), especially with TGF-β1 and -β3 which bind TβRII with high affinity and in turn recruit TβRI by enabling its binding to a composite TGF-β:TβRII interface. This is designated cooperative receptor recruitment and is illustrated in Fig. 1A.

**Figure 1:**
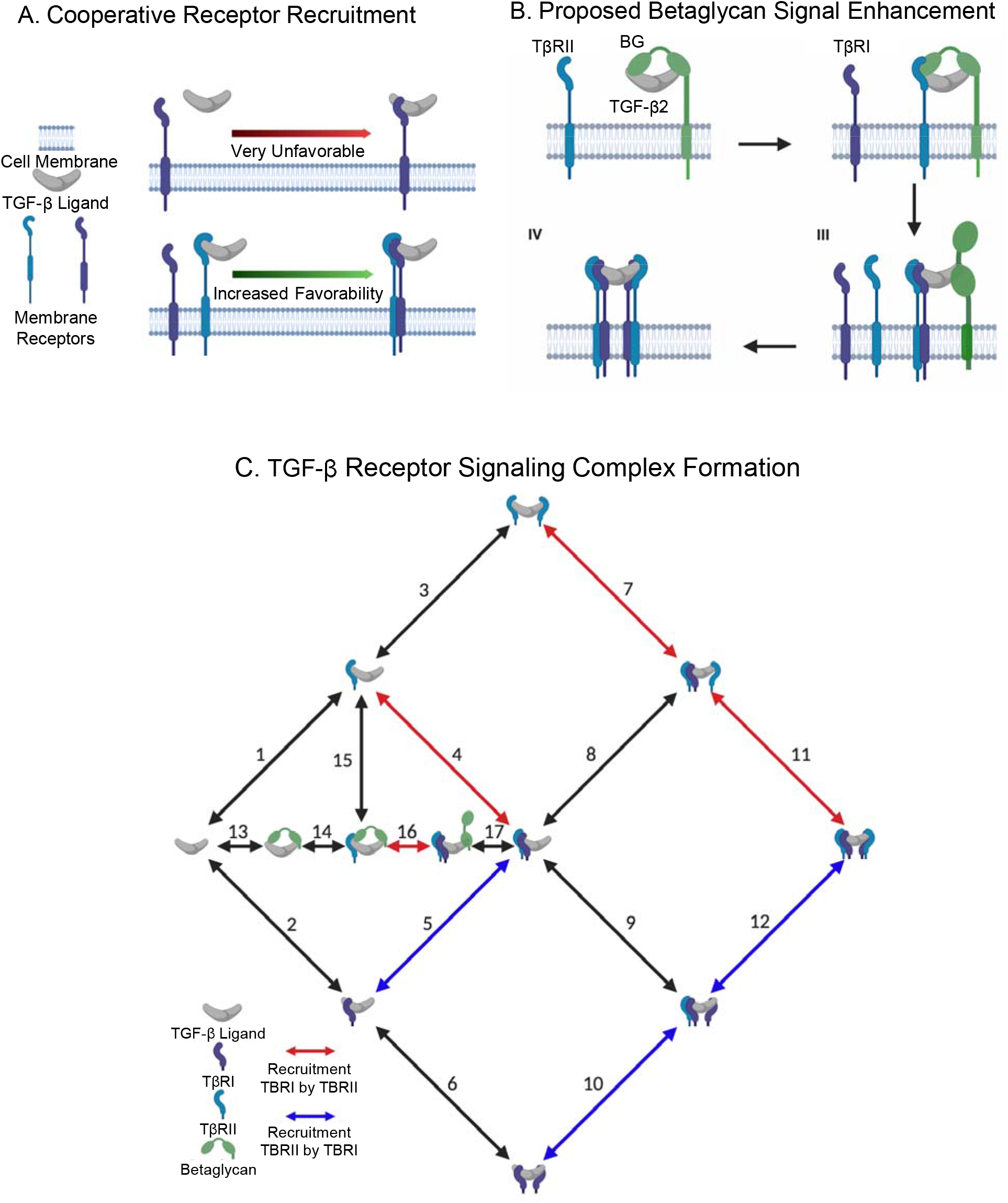
Schematic and biology of TGF-β receptor signaling complex formation. (A) Cooperative receptor recruitment is a biological interaction where the presence of one receptor increases the affinity of another receptor, usually a weak affinity receptor. (B) In the TGF-β2 system, betaglycan with two domains (orphan, BG-O, and zona-pellucida, BG-ZP, domains) is predicted to enhance TGF-β2 signaling by increasing the affinity of the TβRII (blue receptor). Betaglycan dissociates and TβRI (purple receptor) binds, creating half of the tetrameric signaling complex. (C) All three TGF-β ligands (TGF-β1/2/3) signal through a tetrameric signaling complex composed of two Type II receptors (blue bean) and two Type I receptors (purple bean) and its formation can be aided by a membrane bound coreceptor, betaglycan (green bean). Receptor complex assembly is formed through reversible reactions (double-sided arrows). The colored arrows indicate where cooperative receptor recruitment was found and/or tested. The Single-stage Receptor Recruitment (SRR) model accounts for the recruitment of TβRI by TβRII (red arrows) and the Two-stage Receptor Recruitment (TRR) model builds off the SRR model by further incorporating the recruitment of TβRII by TβRI (blue arrows).

The assembly of the complex with TGF-β2, however, is not as well understood as it binds both TβRI and TβRII weakly (9,11,13). Thus, it may not assemble its receptors in the same type II, type I stepwise manner as TGF-β1 and TGF-β3. Moreover, for all three isoforms, although it is conceivable that the second TβRI:TβRII pair assembles in the same manner as the first pair, this is difficult to validate with the experimental tools available today. Computational modeling can shed light on alternative pathways for assembly, both in early and later stages, by assessing how accurately the models recapitulate experimental data for different reaction pathways. Figure 1C reflects this knowledge and depicts the reaction pathways incorporated in our models for assembly of the TGF-β(TβRI)_2_(TβRII)_2_ heterotetramer.

Three deterministic models were constructed to explore the importance of receptor-receptor interaction in TβRI:TβRII heterotetramer assembly. These models were constructed based on published surface plasmon resonance (SPR) data, which provides kinetic rates for many steps in the receptor assembly pathway, though not all. Table S1 shows the rates for the model, including literature references and a brief rationalization for their use.

The Single-stage Receptor Recruitment (SRR) model includes the experimentally proven potentiation of TβRI binding by TβRII (8,11–13), pictured by the red arrows in Figure 1C. The stabilized epitope between TβRII and the ligand for increased binding of TβRI (SRR model) suggests there should also be a similar stabilizing epitope between the ligand and TβRI for increased binding of TβRII. Therefore, the Two-stage Receptor Recruitment (TRR) model was developed, which includes the recruitment represented in the SRR mode, but also a symmetric form of recruitment where ligand-bound TβRI increases the affinity of TβRII to the ligand-complex. Similar symmetric and/or additive cooperative receptor recruitment interactions have been found in other receptor proteins such as RXR nuclear receptor, TCR, Proteinase-activated Receptor2 and TLR4 (29–31). The TRR model is visually represented by the red and blue arrows in Figure 1C. The recruitment of TβRII by TβRI has not been tested with biological methods due to the extremely low affinity of TβRI to the ligand and the resulting difficulty in obtaining the complex for measurement by SPR. The computational approach used in this paper allowed us to test the validity of this symmetric receptor recruitment. In Figure 1C the black arrows are reactions that are the same between all receptor recruitment models developed. The No Receptor Recruitment (NRR) model assumes no potentiation of TβRI by TβRII, nor TβRII by TβRI, and replaces the potentiated rates with kinetic rates for the first addition of a TβRI and TβRII to the ligand. This model acts as a control to measure changes in signaling in the oligomerization pathway.

### BG-mediated potentiation of receptor complex assembly and criteria for model selection

The next component added to our computational models, and the one on which this paper is mainly focused, is the interaction of BG with TGF-β2. The previous data, both the initial cell-based cross-linking studies (17) and the more recent SPR and ITC-based binding studies with purified full-length BG and its subdomains (23,27,32), provide support for a handoff mechanism, as depicted in Fig. 1B. There, however, is still some uncertainty. The available structural and binding data (12,13,27) suggest that a quaternary complex with TGF-β:BG:TβRII:TβRI should exist, yet and in contrast to the TGF-β:BG:TβRII ternary complex, this was not captured in cell-based crosslinking experiments (17). This led to the suggestion that this species was transient, and therefore minimally contributed to the overall signal (27). The computational models created in this paper address the ambiguities of this proposed mechanism for BG-mediated TGF-β2 signaling to identify data-consistent mechanisms and assumptions needed to replicate in vitro BG behavior.

To establish minimal requirements for our BG and receptor signaling simulation, we relied on a combination of quantitative and qualitative BG observations for model evaluation. The evaluation criteria are based on observations from specific experimental results previously published to discriminate BG function. The evaluation criteria for our betaglycan models include: (1) BG increases signal production in TGF-β2 to a greater degree than TGF-β1/3 (15), (2) BG recovers TGF-β2 signaling to levels comparable to TGF-β1/ β3 ligands (15), and (3) BG can act as a dual modulator of TGF-β2 signaling in a concentration dependent manner (33). The third statement in our evaluation criteria has both a strong experimental and theoretical footing and therefore is a reasonable BG behavior to expect. The most notable experimental evidence supporting this statement comes from studies that showed that membrane-bound BG, not the membrane-shed form reported by others (34–36), acts as an antagonist of TGF-β signaling in certain cell lines (33). Theoretically, a co-receptor that sequesters ligand from the extracellular environment to present it to the signaling receptors for binding has the ability to act as a competitive inhibitor at high concentrations. Thus, betaglycan is expected to behave in a biphasic manner, where it potentiates receptor complex assembly and signaling at moderate concentrations, but becomes inhibitory at high concentrations. This type of biphasic effect was demonstrated in 2012 with the BMP coreceptor, CV-2 (37,38), but has also been reported for endoglin, another co-receptor of the TGF-b family that is homologous to the betalycan that promotes the signaling of the TGF-β family ligands, BMP-9 and BMP-10 (39).

### Membrane localization

One of the most important considerations for computational models of this system is to accurately account for membrane localization of the signaling receptors and non-signaling co-receptors through a surface enhancement factor (SEF). The SEF accounts for local increases in concentration and access of interacting receptors, thereby, enhancing second order reactions that occur on the cell membrane. Typically, there are two sequential resistances for a binding reaction between two components: the transport limited step and the reaction limited step. The first advantage of reactions occurring on a surface or a cell membrane, as opposed to in free solution, is that the reactions are essentially two-dimensional after the initial binding event, removing a complete degree of freedom through a process known as reduction of dimensionality. Though diffusional transport of a membrane-bound receptor might be slower relative to free solution, the reduced dimension increases the probability of contacting the partner. This can be modeled by an effective decrease in the dissociation constants and applies equally to all surface-localized reactions (40). Therefore, reactions that take place between two membrane-bound macromolecules will have an increased favorability in comparison to a reaction where an extracellular signaling molecule has to find and favorably orient itself with respect to a transmembrane protein (reactions 1, 2, and 13 in Fig. 1C). Specific reasoning for quantifying the SEF value can be found in the Supplemental Material section S3.

### Simulations that exclude betaglycan

We first analyzed the viability of the three receptor recruitment models (NRR, SRR, TRR) with all three TGF-β ligands, but in the absence of BG. We evaluated the biological relevancy of each model by calculating a root mean square error (RMSE) between models to experimental signaling data in cell lines with no BG (15) (Supplemental Methods). Lower RMSE values indicate the simulated data is more representative of experimental signaling behavior when no BG is present.

The known and unknown parameters, starting conditions that affect the model output, in the no BG simulations were similar across all three ligand types (β1/2/3) and across two of the receptor complex assembly models (NRR and SRR). The known parameters for the first simulations were ligand concentration and receptor concentration. The specific ligand concentrations selected enabled comparison of simulated data to experimental data (15). The starting receptor concentration selected, 160 nM, was the median receptor level found across a wide range of cell lines reported in literature. The values reported as receptors per cell were converted to concentration with volume calculations, as described in Supplemental Material (41) (Supplemental Material). Equimolar concentrations for TβRI and TβRII were used in our simulations based on previous studies (42). To ensure model consistency and integrity, a range of receptor concentrations, 100 to 250 nM, was tested to determine the effect of this parameter on signal performance.

With the biophysical data available in literature, there is only one unknown parameter for each ligand type in the NRR and SRR models. The unknown parameter is the absolute forward and reverse rate constants of TGF-β2 reaction 1 (including homologous reactions, see Supplemental Material), and TGF-β1 and TGF-β3 reaction 2 (including homologous reactions, see Supplemental Material). While the value of the dissociation constant for each reaction is known, the specific forward and reverse rates are not known. To test the impact of these default values on the RMSE calculation, we performed a local sensitivity analysis on the forward and reverse binding steps by simultaneously increasing their values between 1- and 500-fold. The default rates were curated based on similar reactions with measured forward and reverse binding steps and were increased to relevant ranges found in the literature. Simultaneously increasing the fold change in the forward and reverse binding steps preserved the experimentally measured dissociation constant and allowed us to determine the impact of changing the absolute rates on the RMSE calculations. As shown in Figure 2A, increasing the fold change value of the absolute rates to ranges that were observed in literature, did not improve model fit for either the NRR or SRR model across a range of receptor concentrations. Furthermore, decreasing the absolute rate values to different degrees between the three ligand types did not lead to an appreciable change in individual model fitness as measured by the RMSE calculations. For example, in the NRR model, decreasing TGF-β1/3 reaction 2, and other homologous reactions, by 5-fold while decreasing TGF-β2 reaction 1, and other homologous reactions, by 500-fold did not appreciably affect the RMSE analysis (Fig. S1). Due to the minimal impact of changing the absolute rates for both models, the magnitude of the absolute rates chosen were the starting default values (Fig. 2A, blue line). For both models, changing the default receptor concentration of 160 nM minimally affected the signaling pattern as shown in Figure 2A. The results of the local sensitivity analysis for receptor concentration supported the selection of our starting receptor concentration value.

**Figure 2:**
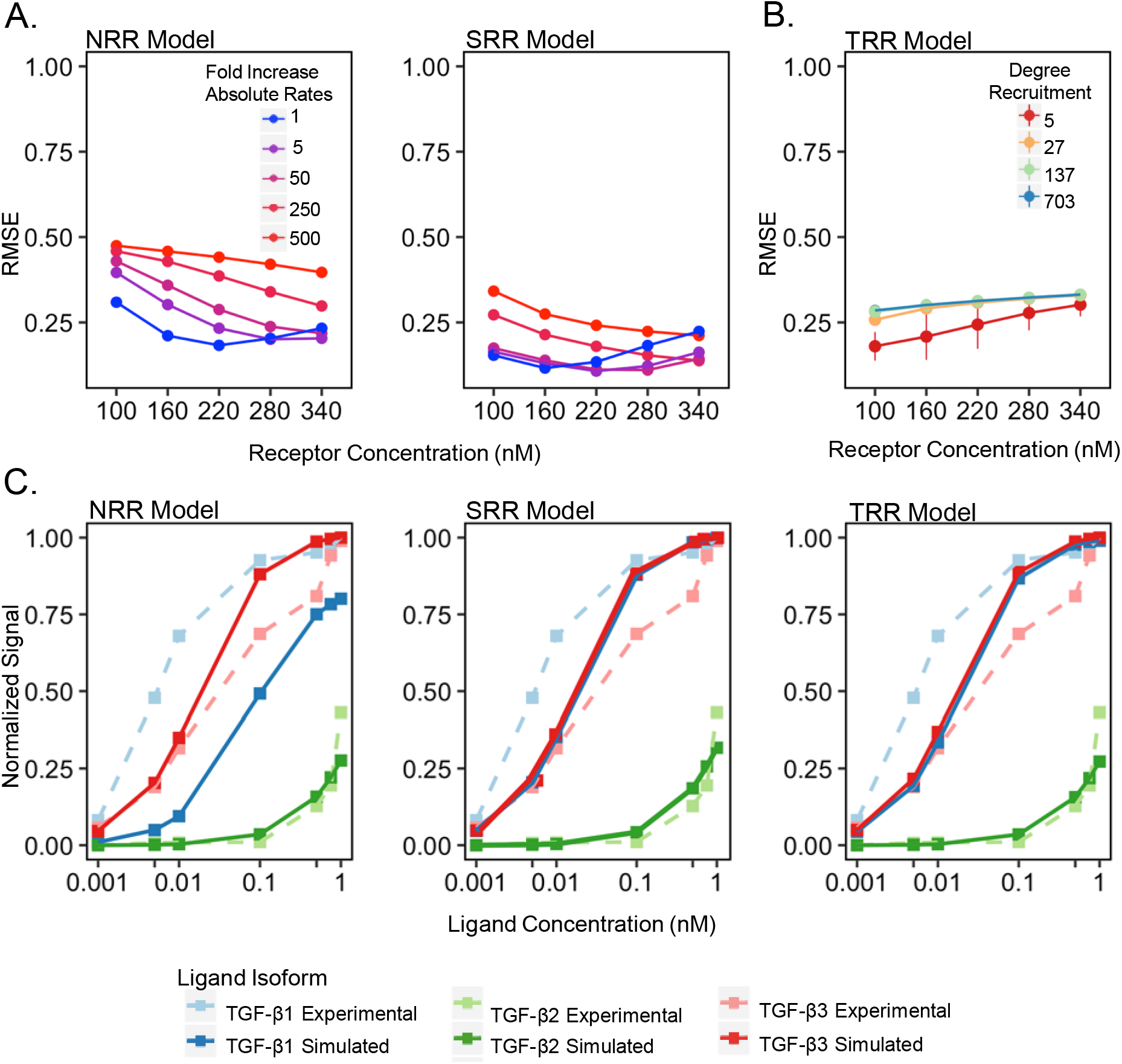
Results of simulated models vs published data and parameter selection when no BG is present. (A) Changing the value of the default receptor concentration, 160 nM, and the default absolute rates (blue line) while having a uniform dissociation constant (on-rate/off-rate ratio), does not improve the RMSE value for the NRR and SRR models. (B) The degree of recruitment (red to blue lines) with the best-fit for the hypothesized receptor recruitment of TβRII by TβRI was approximately 5-fold. (C) With the predetermined SEF value of 50 and receptor concentration of 160 nM across all models, the “best-fit” simulations (solid lines) are able to obtain results similar to experimental data (dotted lines) (Cheifetz et al., 1990).

The second simulation tested the accuracy of the TRR model when no BG was present. The known parameters for this simulation, ligand and receptor concentration, were maintained from the first simulation. The unknown parameters were the degree of receptor recruitment for reactions 5, 10, and 12 (Fig. 1C) and the rates for these reactions. Like the first simulation, decreasing the absolute values of the rates while preserving the dissociation rate constants did not significantly change the RMSE values. Figure 2B shows the effect of increasing the degree of recruitment in the TRR model on the RMSE analysis. The bars at each point represent the minimal impact of changing absolute rates. The degree of recruitment to best fit the experimental data is roughly 5 (Figs. 2B and S3). This value characterizes the degree of increased favorability that a ligand bound TβRI has on recruiting TβRII to the ligand-complex. With the default values used for our unknown parameters in all three receptor recruitment models, there was no appreciable change in the RMSE value when the equimolar assumption for TβRII and TβRI was relaxed (Fig. S4). Therefore, a receptor concentration of 160 nM for TβRI and TβRII was maintained for further computations and should be assumed unless otherwise mentioned.

The parameter sets used for Figure 2C represent a good solution for each model at a receptor concentration of 160 nM, SEF of 50, and default absolute rate values with similar magnitudes to the absolute rates determined in SPR experiments. All three models can produce results that recapitulate TGF-β2 signaling patterns with no BG present, but relative to each other, the SRR and TRR models produce a better fit to the experimental data. TGF-β1 and TGF-β3 were also modeled as a further validation of the working assumptions in each model. Although they are not the focus of our paper, they further validate our simulations by having similar signaling patterns in the SRR and TRR models, as the kinetic rates for each are very similar. As predicted, the NRR model underperforms in reproducing TGF-β1 and TGF-β3 behavior, likely due to the lack of receptor recruitment present in the other models.

### Simulations that include betaglycan

We added BG to the simulations for the second step of modeling TGF-β2 signaling. We evaluated the biological relevancy of each model by calculating a root mean square error (RMSE) between models to experimental signaling data in cell lines with BG and comparing its behavior to the evaluation criteria of BG behaviors reported in the literature and enumerated (15) (Supplemental Methods), but also by comparing to the evaluation criteria of BG behaviors reported in the literature and criteria for model evaluation described above (33,37,38) The results shown in Figure 3A depict how effective BG is in each model at enhancing TGF-β2 signal and shows the percent of tetramer produced out of the signaling species and TGF-β2:BG:TβRII:TβRI. The different points for each model represent the parameter sets across a range of ligand and BG concentrations tested for the three models. A tradeoff is present between the amount of BG induced signal increase and the percent tetramer produced for the SRR and TRR models.

**Figure 3:**
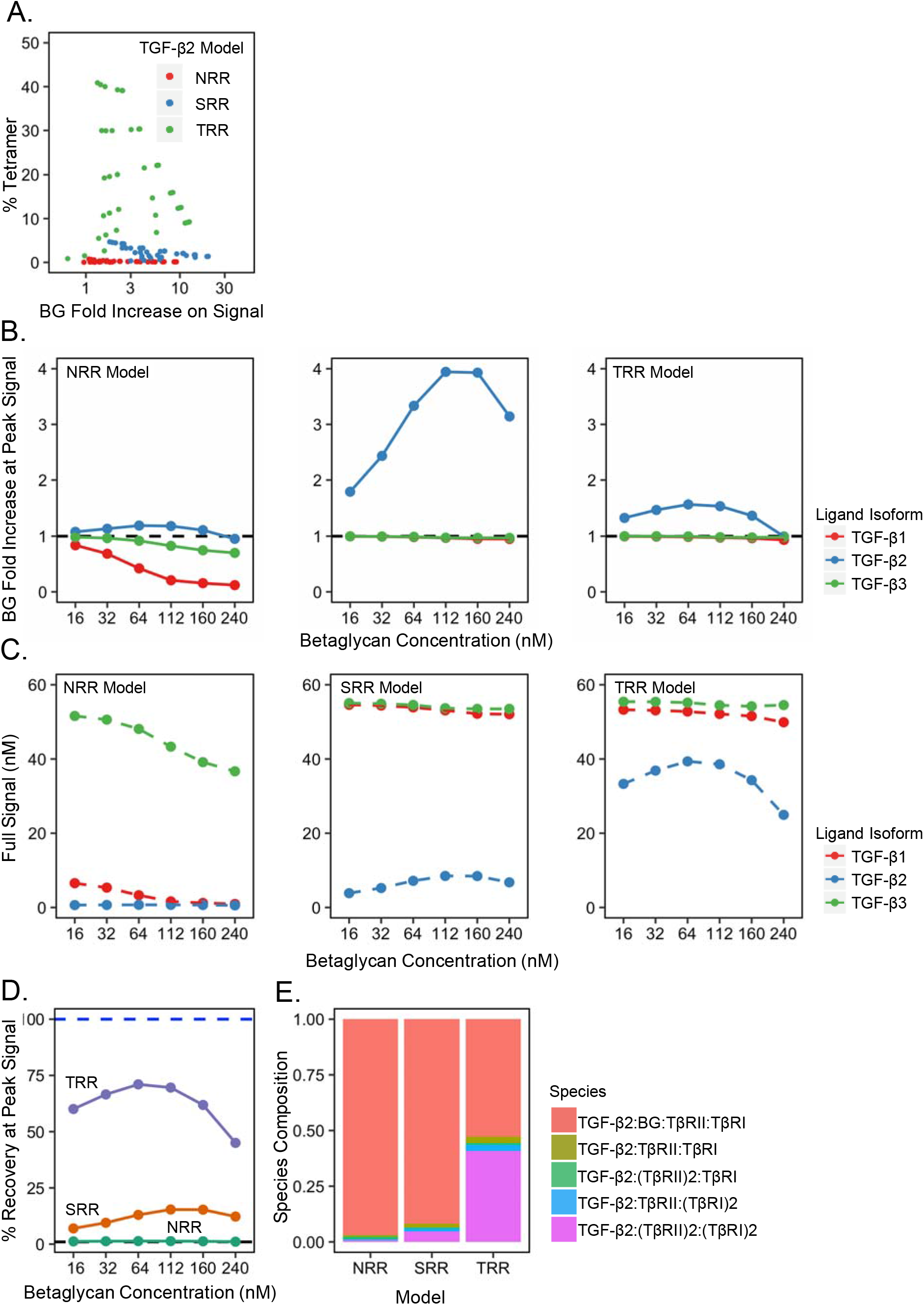
Poor performance in three models when BG is added. (A) The parameter sets with an SEF of 50 and a receptor concentration of 160 nM (colored dots) for each TGF-β2 model are displayed together. A tradeoff between tetramer production and BG enhanced signal is present across all of the models. (B) The signaling enhancement of the three ligand isoforms by BG across the three models was greater in the TGF-β2 (blue line) system than TGF-β1/β3 (green and red lines) systems. The data was normalized to the amount of signal produced in each model with no BG present, therefore, the black dotted horizontal line represents no increase in signal by adding BG. (C) The absolute signaling concentrations (dashed lines) for the three ligand systems (same coloring as Figure 3C) across the three models are shown. The NRR model produced almost no TGF-β2 signal and the TRR model produced two times more TGF-β2 signal than the SRR model. (D) When BG was added (x-axis), the TRR model at 75% recovery, was the best at recapitulating TGF-β2 signal to levels comparable to TGF-β3 signal (blue dashed line). Higher concentrations of BG inhibit TGF-β2 signal as seen by the bell shape curves on the graph. (E) TGF-β2 species composition of the signaling species and TGF-β2/BG/TβRII/TβRI across each model. The TRR model produced more tetramer (bright pink) than the NRR and SRR models, and the major species in all models is TGF-β2/BG/TβRII/TβRI (light red).

The results in Figure 3B show that all three models meet the first and third evaluation criteria of BG behavior: there is a greater positive impact on TGF-β2 signal (blue line) than TGF-β1/3 (red/green lines) and BG can act as a dual modulator of signal in a concentration-dependent manner. The graphs are each normalized to signaling levels with no BG present. The SRR model produces the largest fold change out of the three models due to normalizing technique used, but produces less absolute signal than the TRR model (Fig. 3C). Figure 3D shows how well each model meets the second and third evaluation criterium of BG behaviors: BG produces TGF-β2 signal that is comparable to TGF-β3 signal and BG can act as a dual modulator of signal in a concentration-dependent manner. As demonstrated by a BG concentration of 240 nM, all TGF-β2 models experience a biphasic effect whereby potentiation of signaling by BG is mediated most effectively by intermediate concentrations. The behavior of BG in the TRR model performs the best at potentiating TGF-β2 signal, achieving roughly 75% of TGF-β3 signaling at a BG concentration of about 100 nM. BG, however, provides less then 20% recovery in the SRR model and almost zero percent recovery in the NRR model. A rescue closer to 100 % in TGF-β2 signaling is expected because experimental data shows that BG fully rescues TGF-β2 signal to TGF-β1 and TGF-β3 signaling levels (15).

To investigate the causes of suboptimal TGF-β2 signal rescue, we examined the concentrations of all species at steady-state to obtain a better understanding of the behavior of our models. Figure 3E is a breakdown of the signaling species and TGF-β:BG:TβRII:TβRI composition at steady state for each model. None of the models produce greater than 50% TβRI:TβRII heterotetramer (bright pink) and the major species in all of the models is TGF-β:BG:TβRII:TβRI. This is at odds with the proposal that the TGF-β:BG:TβRII:TβRI complex is transient, thus lowly populated (27). This inconsistency may on the one hand indicate that the TGF-β:BG:TβRII:TβRI complex accumulates on the cell membrane and is not transient as previously thought. If this species is not transient, then allowing this species to contribute to signal may improve the suboptimal signal recovery. It is possible on the other hand that the kinetic rates for the dissociation of the TGF-β:BG:TβRII:TβRI complex inaccurately capture the predicted transience of this intermediate species. Since no kinetic rates were reported for disassociation of BG from the TGF-β:BG:TβRII:TβRI complex, the starting kinetic rate for the dissociation of the BG quaternary species was the dissociation of the BG-ZP domain from free ligand. This inferred dissociation constant may inaccurately represent the transience of the BG quaternary species. Due to the ambiguity surrounding the transience and signaling capability of the BG quaternary species, the assumption that this species is transient was tested with the following sections.

### Testing assumption: BG quaternary species is transient

Previously, we did not allow the TGF-β:BG:TβRII:TβRI complex to signal based on the assumption that it was a transient species. If we no longer apply the transient assumption to our models then the TGF-β:BG:TβRII:TβRI complex (Fig. 4A, species circled in red) would be able to contribute to overall signal based on the earlier studies which showed that TβRI:TβRII heterodimers are capable of signaling (8).

**Figure 4:**
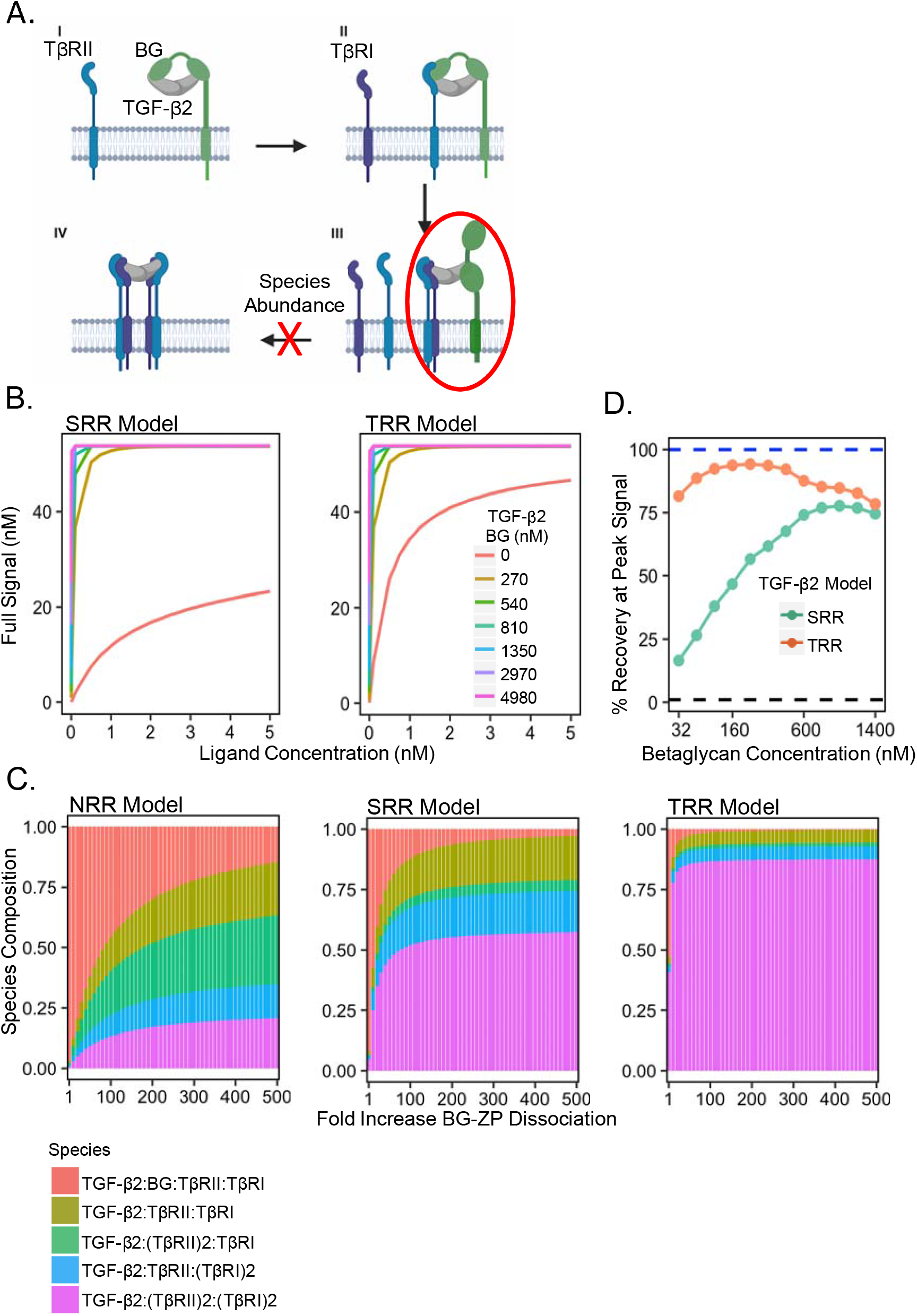
TGF-β2/BG/TβRII/TβRI inconsistency solved by small increase in BG-ZP dissociation to ligand-complex. TRR model outperforms other models in recapitulating betaglycan behavior. (A) A depiction of the inconsistency previously found where TGF-β2/BG/TβRII/TβRI complex (circled in red) was the most prevalent species out of all the species created. This abundance was hypothesized to inhibit the formation of signaling species. (B) When TGF-β2/BG/TβRII/TβRI species is not required to be transient and contributes to signal, there is no biphasic effect across a wide range of ligand concentrations (x-axis) and BG concentrations (colored lines), 0.001 to 5 nM and 0 to 4980 nM, respectively. (C) A species composition analysis at peak signal, with a receptor concentration of 160 nM, and a SEF of 50, shows minimal increases in BG-ZP dissociation reduces the abundance of TGF-β2/BG/TβRII/TβRI species in the SRR and TRR models. NRR model requires a larger increase in BG-ZP dissociation compared to SRR and NRR models. The BG-ZP dissociation fold increase selected for each model was when increasing the BG-ZP dissociation did not impacted the signaling results with the parameter ranges tested. This behavior is reached whenever 90% of the maximum signal is achieved. (D) TGF-β2 signal in the TRR model (orange line) can recover greater than 95% of TGF-β3 signal (blue dashed line) while the TGF-β2 signal in the SRR model (green line) can recover roughly 75% of TGF-β3 signal.

When we allowed this species to contribute to signal, we observed across an extremely wide combination of ligand concentrations and BG concentrations that BG could not act as a dual modulator of signal in a concentration-dependent manner (Fig. 4B) since BG no longer had an optimal ratio of BG to signaling receptors for TGF-β2 signal potentiation. These results demonstrate if TGF-β2:BG:TβRII:TβRI is not a transient species it should not contribute to signal in order to recapitulate the ability of BG to promote and reduce TGF-β2 signaling in a concentration-dependent manner. If the transient hypothesis is true, then the inconsistent accumulation of TGF-β:BG:TβRII:TβRI species may be a reason for the suboptimal TGF-β2 signal recovery. A screen for the dissociation of BG-ZP domain in the TGF-β:BG:TβRII:TβRI species (reaction 17) was performed to test if the starting SPR kinetics are a reason for the impeded rescue.

### Effects of increasing BG-ZP dissociation

To further investigate how the two BG domains work together to enhance TGF-β2 signal recovery, we increased the dissociation of the BG-ZP domain between a range of 1-500. This allowed us to determine if reducing the abundance of TGF-β2:BG:TβRII:TβRI (Fig. 4A, species circled in red) could lead to the formation of more TβRI:TβRII heterotetramer, and therefore, increase BG potentiation of TGF-β2 signaling.

The NRR model can produce results where the TGF-β2:BG:TβRII:TβRI species is less than 50% of the signaling species, but this requires 100-fold or greater increases in BG-ZP dissociation (Fig. 4C). The SRR and TRR models, in contrast, produce significantly less TGF-β2:BG:TβRII:TβRI and abundant TβRI:TβRII heterotetramer, even with modest increases in BG-ZP dissociation (Fig. 4B). Compared to the SRR and TRR models, the NRR model performs poorly at reducing the abundance of TGF-β2:BG:TβRII:TβRI species. The NRR model was therefore left out of further analyses due to its substandard performance and for ease of comparison between the SRR and TRR models. For each model, we selected a fold increase in BG-ZP dissociation which produced at least 90% of the maximum signal for each model (100- and 30-fold increase for the SRR and TRR models, respectively). Selecting a fold increase in BG-ZP dissociation beyond the selected value minimally affects the signaling results of each model. This idea is visualized by the logarithmic shaped curve in species composition graph as the fold change in BG-ZP dissociation increases (Fig. 4C).

Minimally increasing the BG-ZP dissociation in the SRR and TRR models led to a higher percent recovery in TGF-β2 signal. Using the same logic in Figure 3D, Figure 4D shows the TRR model can now achieve greater than 95% recovery of peak signal produced by TGF-β3 while the SRR model produces roughly 75% with a wide range of BG concentrations tested (32 nM to 1400 nM). Comparing Figure 3D and Figure 4D, the signal recovery levels are higher for both the SRR and TRR models when TGF-β2:BG:TβRII:TβRI is predicted to be transient and the dissociation of the species is minimally increased. Thus, these results suggest that BG-ZP quickly dissociates when TβRI is bound and the TGF-β2:BG:TβRII:TβRI species does not signal or has a minor contribution to the overall signal. To test this experimentally, it will be important to further measure the impact of BG on TGF-β signaling under various amounts of BG overexpression.

The mechanism proposed by Villarreal (27) with published rates from SPR experiments does not meet our evaluation criteria if the TGF-β2:BG:TβRII:TβRI species is allowed to contribute to signal because BG no longer acts as a dual modulator of signal, regardless of concentration. However, if the disassociation of TGF-β2:BG:TβRII:TβRI complex is increased, and thus the complex becomes more transient, our results demonstrate greater TGF-β2 signal rescue. Comparing two other system behaviors, percent tetramer and BG-induced signal enhancement, the models that include an increase in BG-ZP dissociation from the ligand-receptor complex outperform models with the starting SPR data (Fig. 5A). Figure 5B summarizes the effect of increasing BG-ZP dissociation on model performance at varying concentrations of BG (red to blue lines). This RMSE analysis not only compares the signaling differences between TGF-β2 with BG to TGF-β3 without BG, but also incorporates the expected signaling behavior of TGF-β1 and β3 systems with and without BG. The TRR model once again outperforms the SRR model at recapitulating the BG behaviors. Although the SRR model is supported by SPR measurements, it is nonetheless possible that other forms of cooperative receptor recruitment are present in the TGF-β system (8,12,13). A symmetric form of recruitment, as exemplified by the TRR model, better facilitates the formation of the heterotetramer which follows the pattern in other protein systems (29). The TRR model also outperforms the SRR model on almost all evaluation criteria of BG behavior. The simulations presented therefore suggest that the TRR model is the most realistic model tested. To enable experimentally testing of the conclusions presented in this paper, further differences between the SRR and TRR models were investigated, as summarized in the section that follows.

**Figure 5:**
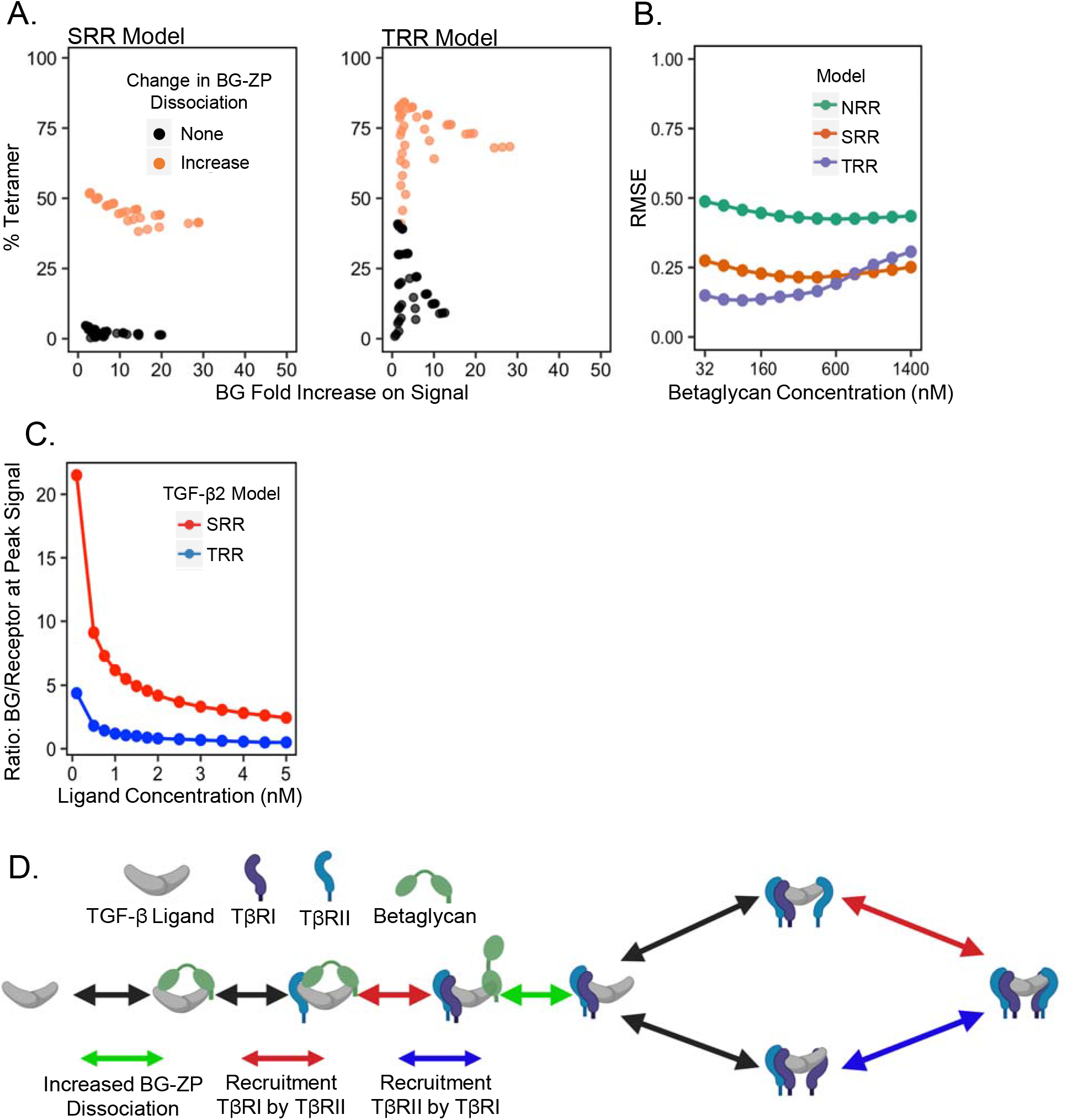
Distinguishing between models and BG-mediated TGF-β2 signal hypotheses. (A) Across a broad range of parameter sets, almost any increase in the dissociation of BG-ZP (orange points) outperforms the original parameter sets with no increase in BG-ZP dissociation (black points). (B) When requiring the TGF-β2/BG/TβRII/TβRI species to be transient, the RMSE analysis captures how the models measure up to no BG and BG system requirements in all three ligand systems. The NRR model was incorporated again to show effectiveness of adding cooperative receptor recruitment into the TGF-β receptor signaling complex formation in the presence and absence of BG. Each model shows BG’s inhibitory effect on TGF-β’s signal by the concave shape of the graphs. Across a wide range of BG concentrations (x-axis) the TRR model recapitulates no BG and BG behavior the best in all three ligand systems until BG inhibits TGF-β signal causing the RMSE to increase. (C) A testable distinction between the SRR and TRR models was found in the BG to receptor ratio required to achieve the biphasic effect on TGF-β2 signal by BG. Across a wide range of ligand concentrations, 0.001-5 nM, the SRR model (red points) needs 4.7-4.9 time more BG to induce the biphasic effect than the TRR model (blue points). (D) Zoomed in diagram of the receptor complex assembly that highlights the final conclusions. The red and blue arrows represent the types of cooperative receptor recruitment that is hypothesized to be present and the green arrow indicates an increase in the dissociation of TGF-β2/BG/TβRII/TβRI improves model performance under certain conditions.

### Investigating model differences

One obvious difference between the SRR and TRR models is the varying betaglycan concentrations needed to induce an inhibitory effect on signal (Fig. 4D). Therefore, we sought to determine the BG to signaling receptor (TβRI and TβRII) ratio that is required to produce a biphasic effect under the assumption that TGF-β2:BG:TβRII:TβRI is transient with a model specific increase in the BG-ZP dissociation (Fig. 4B). With a similar ligand concentration in each system, the SRR model needed a 2.44 to 21.5 BG to TβRI/TβRII ratio to induce a biphasic effect, whereas the TRR model required a 0.50 to 4.38 BG to TβRI/TβRII ratio to induce a biphasic effect (Figure 5E). Thus, the SRR model needed a 4.7-4.9 times greater concentration of BG to receptor ratio than the TRR model. Experiments that seek to identify the molar ratios of receptors and betaglycan will provide useful data to discriminate between these alternatives.

## Discussion

The role of BG in selectively facilitating TGF-β2 signaling has been heavily investigated, but the mechanism is still not fully understood. In the absence of BG, physiological levels of TGF-β2 are insufficient to produce efficient signal to initiate targeted gene expression, leading to a disruption of developmental processes (45).The focus of this work was to identify conditions for selective enhancement of TGF-β2 signaling and provide support for additional receptor binding cooperativity that has been difficult to test with experimental tools currently available. Through modeling of the kinetic mechanism, it appears that BG binding to TGF-β2 ligand through two domains effectively potentiates TGF-β2 signal and a symmetric cooperative receptor recruitment between TβRI and TβRII best explains the experimental data (TRR model).

Computational modeling demonstrated that the proposed BG-mediated potentiation of TGF-β2 signaling with inferred SPR rates produced suboptimal signal rescue, with none of the models producing more than than 75% signal recovery. This indicated that the transient hypothesis of the TGF-β2:BG:TβRII:TβRI species is correct, or there may be a more complex biological interaction taking place between the macromolecules that the inferred SPR data could not accurately represent. If the TGF-β2:BG:TβRII:TβRI species is allowed to signal, TGF-β2 signal recovery is increased, but there is no biphasic effect across a wide range of BG concentrations. Due to the inability for BG to act as a dual modulator of signal, we conclude that BG quaternary species is likely transient, and thus, its overall contribution to signaling is limited.

When the dissociation of the TGF-β2:BG:TβRII:TβRI species was increased, through an increase in the BG-ZP dissociation rate constant, the TRR model reached about 95% rescue of the TGF-β2 signal, while the SRR model reached approximately 75% rescue. The improvement in TGF-β2 signal rescue from minimal increases in BG-ZP dissociation in the TRR model highlights the importance of this step to predicting model performance. With these findings, it is possible that the binding of TβRI to the TGF-β:BG:TβRII complex increases the dissociation of the BG-ZP domain through steric interactions or a conformational change. This interaction cannot be easily measured in real time with experimental tools available today, but our models identified the importance of this dissociation constant in determining model performance. These results support the hypothesis that there may be a more complex biological interaction taking place at this reaction than originally inferred.

The TRR model with a minor increase in BG-ZP dissociation, meets all TGF-β predicted behavior with and without BG present. The differences in the performance of the TRR, SRR, and NRR models indicate the importance of receptor-receptor interactions throughout the TβRI:TβRII heterotetramer assembly pathway. Our modeling results favored the TRR model no matter the alteration in the BG mechanism. TRR’s robust performance in meeting experimental data, suggests that increasing TβRI’s affinity in later stages of TβRI:TβRII heterotetramer assembly greatly affect the performance of TGF-β signaling behavior. Furthermore, the order of addition for the second pair of TβRI:TβRII to the TGF-β homodimer may be nonspecific due to the increase in TGF-β signaling performance following a minor increase in the binding affinity of TβRI downstream. If the TRR model is operative, then the BG to receptor concentration ratio will be from 0.5 to 4.38 whereas the SRR model has a BG to receptor ratio of 2.44 to 21.5. This is a testable difference between the two systems that could be performed to determine if TβRI does recruit TβRII. Modeling alone does not disprove or prove a model but suggests the TRR model should be further tested to determine estimated quantities relative to receptors in the system.

Even though BG’s biphasic effect on TGF-β signaling is heavily supported, no direct experiments have been performed to show BG can act as a dual modulator of signal in a concentration-dependent manner. To test this statement in our evaluation criteria, BG can be titrated into a cell culture to determine if there is an optimal ratio of BG to signaling receptors that potentiates TGF-β2 signaling activity.

Future work can be performed with stochastic simulations to determine why TGF-β2 evolved to require BG for efficient signaling. Identifying if there are differences in noise and/ or transmission properties in the signaling dynamics between the three ligand systems could highlight potential signaling advantages in a system that requires a co-receptor for proper signaling.

## Materials and Methods

### Computational Tools

We carried out deterministic modeling using a python ODE solver program called pySB. PySB is a framework for building mathematical rule-based models of biochemical systems (43). The deterministic model calculates a concentration of each individual species in simulation under different conditions. The nuclear pSmad signal from each TGF-β signaling species was calculated using a computational model of intracellular TGF-β signaling adapted from Schmierer et al., 2008 (44). The full pSmad signal used in RMSE analysis was calculated from a weighted sum of TGF-β signaling species. The detailed equation can be found in the Supplemental Material. PySB codes for the three receptor recruitment models can be found through GitHub (https://github.com/ingle0/Thesis-Code).

## Figure Legends

**Supplemental Figure 1:**
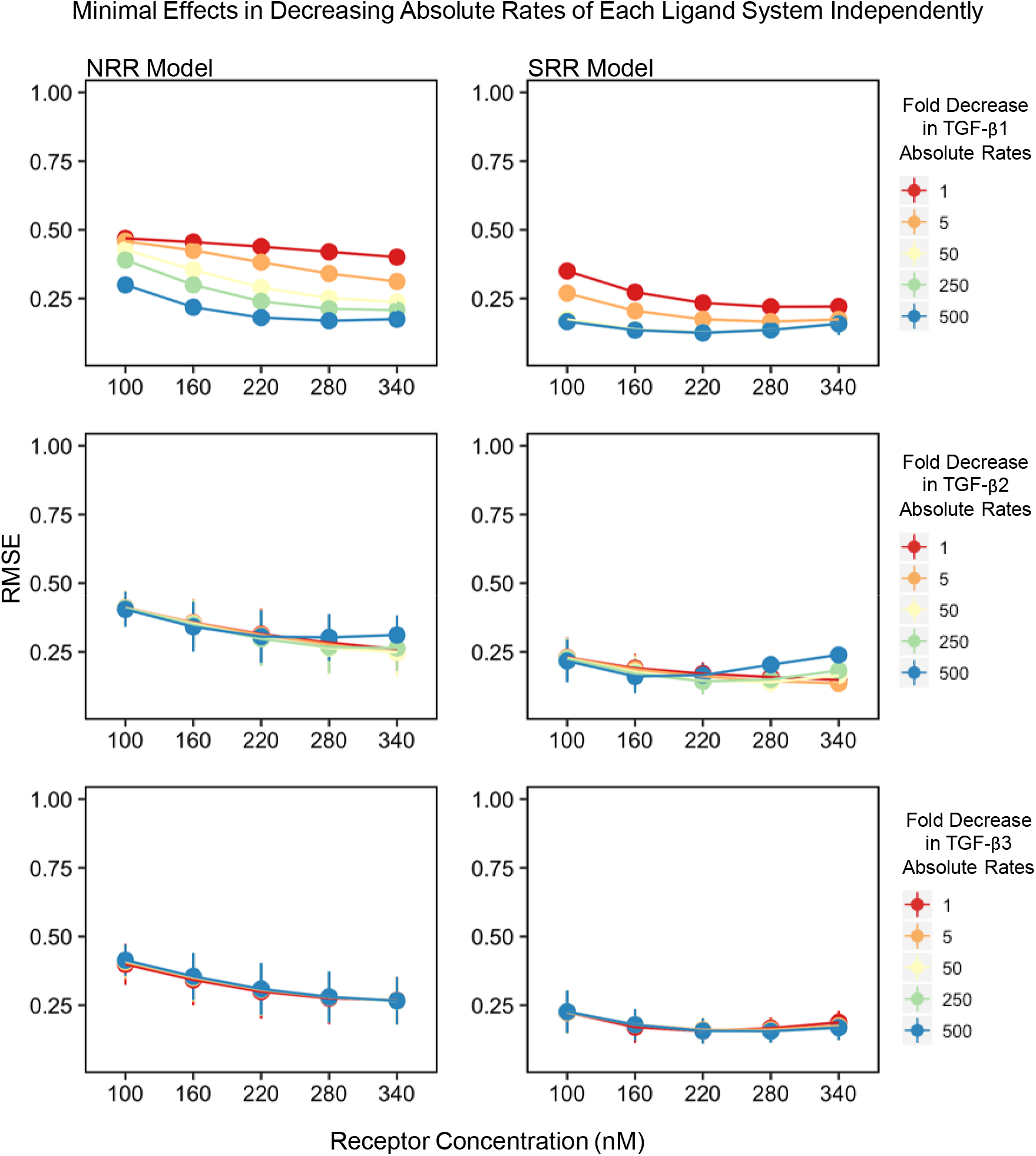
There are minimal effects in decreasing the absolute rates of each ligand system independently. (A) In both the NRR and SRR models, when the fold decrease in TGF-β1 rates are held constant (red to blue lines) and TGF-β2 and TGF-β3 absolute rates are varied (bars present at each dot) there is minimal change in the RMSE value. This pattern remains constant across a range of receptor concentrations (x-axis) from 100 to 340 nM. (B) When the fold decrease in TGF-β2 rates are held constant (red to blue lines) and TGF-β1 and TGF-β3 absolute rates are varied (bars present at each dot) there is minimal change in the RMSE value. This pattern remains constant across a range of receptor concentrations (x-axis) from 100 to 340 nM. (C) When the fold decrease in TGF-β3 rates are held constant (red to blue lines) and TGF-β1 and TGF-β2 absolute rates are varied (bars present at each dot) there is minimal change in the RMSE value. This pattern remains constant across a range of receptor concentrations (x-axis) from 100 to 340 nM.

**Supplemental Figure 2:**
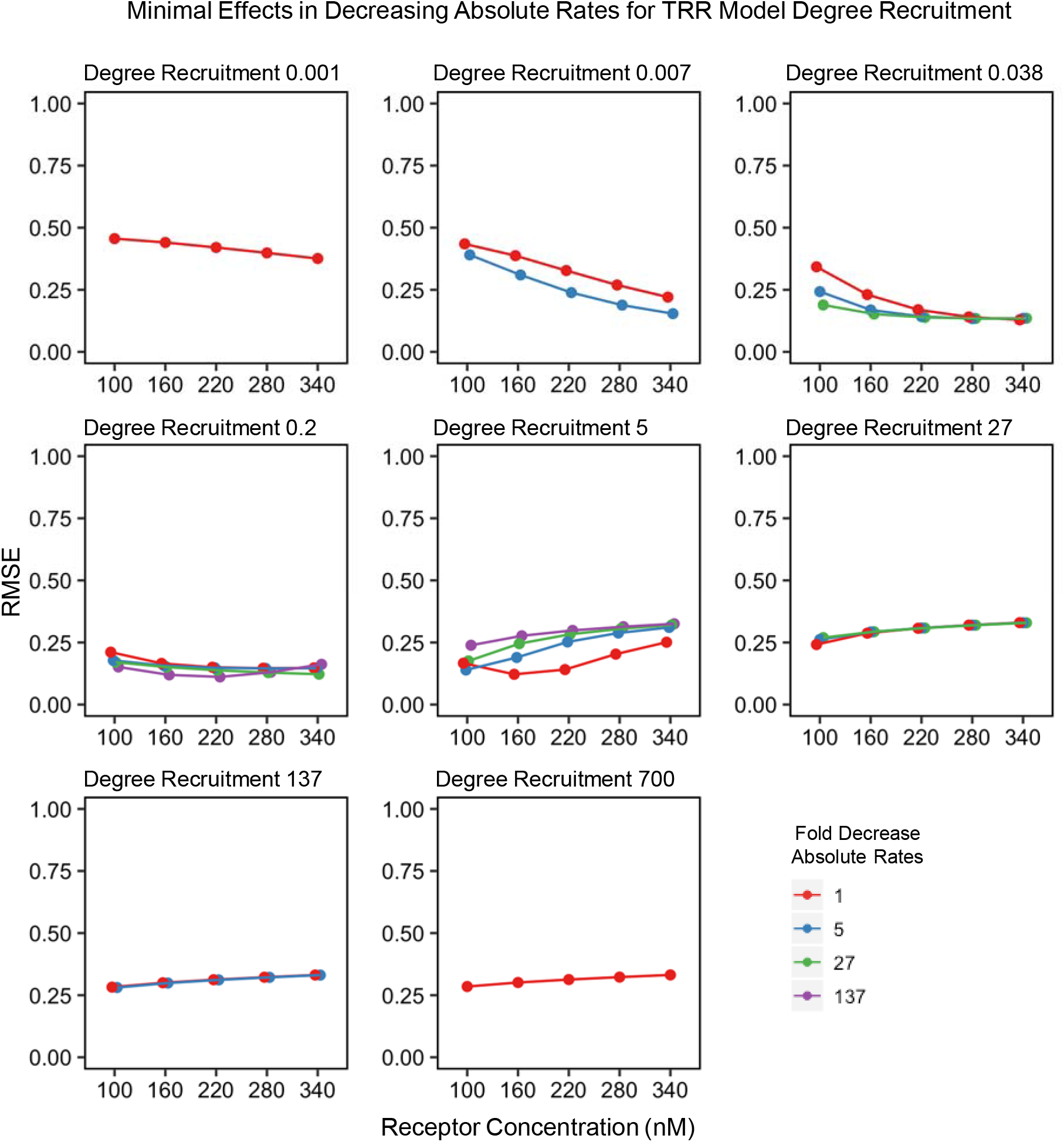
Minimal impact in the RMSE value when decreasing the absolute rates (red, blue, green, and purple lines) across multiple values for TBRI’s recruitment of TBRII. Each graph represents one value for the degree of recruitment TBRI has on TBRII across a range of receptor concentrations (x-axis). Due to the computational screen set up, the median values of degree recruitment have more absolute rates tested than the end values. Degree recruitment of 0.001 and 700 has only one absolute rate (1-fold decrease) shown. Degree of recruitment 0.007 and 137 have two absolute rate values (1 and 5-fold decrease) shown. Degree of recruitment 0.038 and 27 have three absolute rate values (1, 5, and 27-fold decrease) shown. Degree of recruitment 0.2 and 5 have four absolute rate values (1, 5, 27, 137-fold decrease) shown.

**Supplemental Figure 3:**
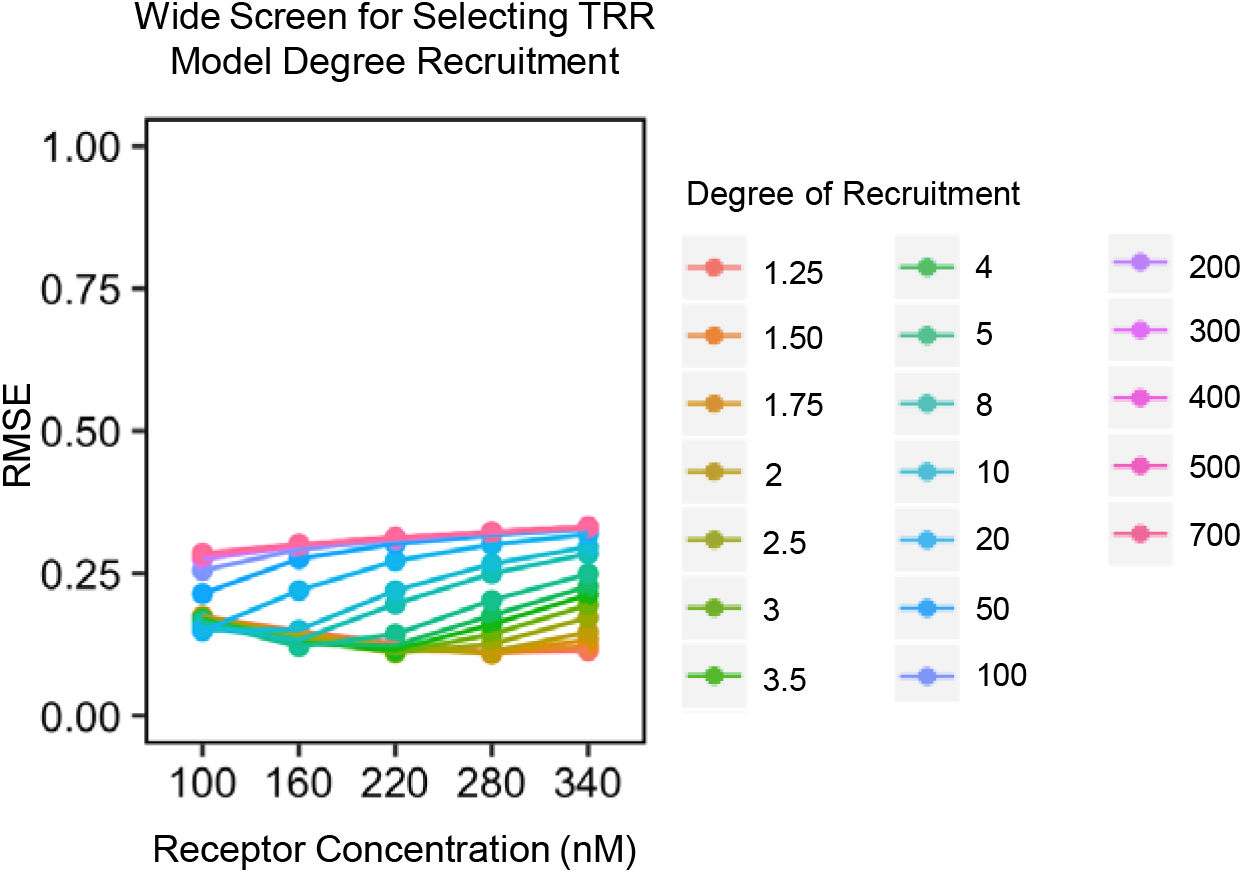
Across a broad range of values for the degree of recruitment TBRI has on TBRII (colored lines), a 5-fold degree recruitment produced the lowest RMSE at a receptor concentration of 160 nM.

**Supplemental Figure 4:**
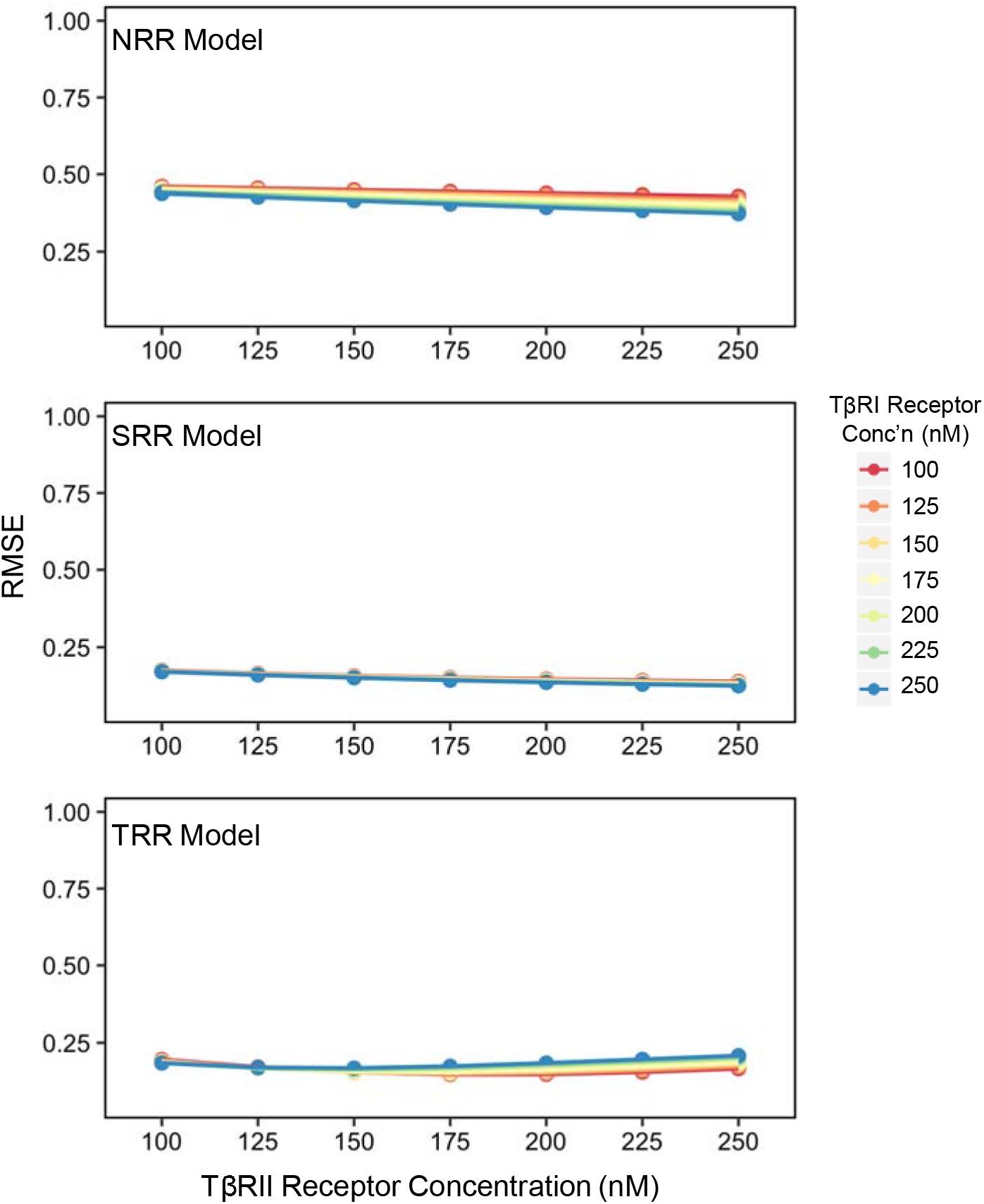
With no BG present, an SEF equal to 50, and specific absolute rates chosen, altering the concentrations of TβRI (red to blue lines) and TβRII (x-axis) independently, minimally affect the RMSE analysis (y-axis).

