## Supplementlal materials for "Computational modeling of TGF-β2:TβRI:TβRII receptor complex assembly as mediated by the TGF-β co-receptor betaglycan"

*S1. Root mean square error analysis*

Root mean square error (RMSE) measures model fit or how accurately the simulated model predicts the experimentally determined biological response. RMSE is the standard deviation of the unexplained variance between the models simulated output and the experimental data (Equation 1).

Equation 1:

$$RMSE=\sqrt{\frac{\sum_{i=1}^{n} ({{Simulated}_{i}-{Experimental}_{i})}^{2}}{n}}$$

Lower RMSE values indicate a better model fit, or less unexplained variance between the simulated and experimental data points. This analysis indicates the absolute fit of the simulated models created to the experimental data found in Cheifetz et al., 1990. Experimental data was extracted from the graph using WebPlotDigitzer (Drevan et al., 2016).

*S2. Creating the models*

We developed a deterministic model of the TGF-β receptor complex assembly that incorporated on-rates and off-rates for each reaction listed in Figure 1A and Supplemental Tables 1-9. Surface plasmon resonance (SPR) data found in literature was the biophysical data used as the starting point for each of the model’s kinetic values. Supplemental Tables 1-9 show the kinetic rates used for each of the three models and their source (No Receptor Recruitment, Single-stage Recruitment, and Two-stage Recruitment).

TGF-β’s signal transduction pathway activates when one type II receptor and one type I receptor bind and continues to build towards full signaling capacity when a heterotetrameric complex of two type II (TΒRII) and two type I (TΒRI) receptors is formed on the cell membrane. The heterotetrameric signaling complex (TGF-β/ TβRII/TβRII/TβRI/TβRI) has about four times the amount of nuclear pSmad accumulation than the dimeric signaling complexes (TGF-β/TβRII/TβRII/TβRI, TGF-β/TβRII/TβRI/TβRI, TGF-β/TβRII/TβRI) (Huang et al., 2011). This was reflected in the model by calculating a total receptor signal that included the full concentration of the TGF-β/TβRII/TβRII/TβRI/TβRI plus ¼ the concentration of the TGF-β/TβRII/TβRI, TGF-β/TβRII/TβRII/TβRI, and TGF-β/TβRII/TβRI/TβRI, otherwise referred to as full signal (Equation 2).

Equation 2:

$$Full Signal=\left[ TGF-\beta\bullet T\beta RII\bullet T\beta RII\bullet T\beta RI\bullet T\beta RI \right]+\frac{1}{4}\left[ TGF-\beta\bullet T\beta RII\bullet T\beta RI \right]+\frac{1}{4}\left[ TGF-\beta\bullet T\beta RII\bullet T\beta RII\bullet T\beta RI \right]+\frac{1}{4}[TGF-\beta\bullet T\beta RII\bullet T\beta RI\bullet T\beta RI]$$

*S3. Surface enhancement factor justification*

An important consideration in modeling any receptor complex assembly is considering the differences in reaction affinities between reactions that involve extracellular and cellular reactants (ligand plus membrane bound receptor) and reactions between two cellular reactants (ligand-receptor complex plus a membrane bound receptor). The magnitude of these reaction affinity differences, defined as surface enhancement factor (SEF), can be challenging to quantify and may seem arbitrary. However, when we examined the relative impact of high and low SEF values and checked with prior literature, we were confident that our SEF choice was valid. If the SEF value is too high it will improve the favorability of every reaction to a degree that washes out distinguishable signaling patterns between models or diminishes the appropriate effect of unfavorable reactions. If it is not applied to the system or is too low, the model will not produce enough signal to accurately fit the simulation data to experimental data. It would be unrealistic to say there is an exact number to fit this interaction, but a value of 50 has been used in previous papers with similar quantitative biological models which have been experimentally validated (Karim et al., 2012; Schmierer et al., 2008). Therefore, a value of 50 will be the baseline for our computational experiments. The No Receptor Recruitment model can also be used to partially validate the SEF selected. If the simulation results from the No Receptor Recruitment model looked exactly like the other two models, the SEF selected may be too high or low as it would wash out the important effects of altering certain reactions. This is not the case in the models presented.

In all the models and ligand systems, the SEF was applied to reactions 3 through 12 and reactions 14 through 17 and were modeled as second order reactions due to both reactants being located on the cell membrane. The SEF was not applied to reactions 1, 2 and 13 due to one of the reactants, the TGF-β ligand, existing in the outside environment instead of on the cell membrane. Due to the creation of the products from a free-floating molecule and a membrane bound receptor, these reactions were modeled as pseudo-first order reactions.

*S4. Homologous reactions*

The SPR data used for the reaction kinetics can only measure interactions between molecules/complexes it can isolate. Due to the unfavorable state of some complexes as well as their complex interactions, analyzing the reactions at these higher order intermediate states, like TGF-β/TβRII/TβRI/TβRI reacting with TβRII, is not yet possible. Therefore, we estimate these reactions by assuming they are homologous to lower order reactions that do have SPR data. Justification for homologous reactions are different for each of the receptor recruitment models and are further explained in the sections below describing the creation of each model in detail.

*S5. No Receptor Recruitment model creation*

The No Receptor Recruitment (NRR) model was created as a control model to ensure that the receptor recruitment applied to SRR and TRR models was indeed required for effective signaling that met biologically known behaviors of the TGF-β system. The difference in signaling patterns between the NRR model and the SRR and TRR models also helps validate the SEF selected.

The justifications for the homologous reactions are the same across the three ligand systems for the NRR model, but the biophysical data is not (Supplemental Tables 1-3). Reactions 3, 5, 8, 10, and 12 are homologous to reaction 1 because there is an addition of a single TβRII to the ligand complex. Therefore, they will have the same dissociation, on-rate, and off-rate constants. Reactions 4, 6, 7, 9, and 11 are homologous to reaction 2 because there is an addition of a single TβRI to the ligand complex. Therefore, they will have the same dissociation, on-rate, and off-rate constants. Reaction 15 is homologous to reaction 13 because there is an addition of BG to the ligand complex.

Reactions 13 through 17 describe the interactions with BG involved in each of the ligand systems. Due to the limited biophysical data of BG interaction with TGF-β1 and TGF-β3, the rates and justifications for reactions 13, 14, 15, and 17 in the TGF-β2 system were maintained. With the same kinetics applied across all three ligand systems we are able to test if our models are robust to the knowledge that BG has an insignificant effect on TGF-β1 and β3 signaling especially in comparison to TGF-β2. In the reactions 13 through 17, reaction 16 is the only rate that changed between the three ligand systems due to the homology to reaction 2 as mentioned previously.

Reaction 13’s dissociation constant (K_D_), on-rate, and off-rate values were taken from Hinck’s 2018 paper. Reaction 14’s dissociation constant was taken from Villarreal et al., 2016. When BG is bound to TGF-β2 it increases the affinity for TβRII. Through SPR data, reaction 14 is most similar to a type II receptors affinity for the TGF-β3 ligand without BG present (Villarreal et al., 2016; Radaev et al., 2010). Therefore, the off-rate used for reaction 14 was from the addition of TβRII to TGF-β3 in the TGF-β3 system (Radaev et al., 2010: Table S2). The off-rate was then divided by the dissociation constant to find the on-rate. Reaction 17 is the addition of BG’s zona pellucida domain to the ligand. The K_D_, on-rate, and off-rate of this reaction were found in Kim et al., 2019. The reaction is represented as a second order reaction and is coded into the system as seen in the following tables to keep the kinetic integrity of the reactions.

*Remaining reaction justifications for NRR model of TGF-β1*

Reaction biophysical data available and selected for the No Receptor Recruitment model of TGF-β1 can be found in Supplemental Table 1. The dissociation constant, on and off-rates for reaction 1 were selected from Huang et al., 2014 publication based off of expertise knowledge as well as the latest and most abundant biophysical data available. The dissociation constant for reaction 2 was found in Radaev et al., 2010. Since the absolute rates (on and off-rates) were not published with the dissociation constant, a screen was run in order to determine if the uncertainty in the absolute rates needed further consideration and testing in our model. As previously demonstrated in Figure 2A, S1, and S3, changing the absolute rates did not significantly affect the results of the system, so rates close to the magnitude of previously observed SPR experiments were selected (Supplemental Table 1).

*Remaining reaction justifications for NRR model of TGF-β2*

Reaction biophysical data available and selected for the No Receptor Recruitment model of TGF-β2 can be found in Supplemental Table 2. The dissociation constant for reaction 1 was chosen by latest published rate and expertise advice, Villarreal et al., 2016. Since the absolute rates (on and off-rates) were not published with the dissociation constant, a screen was run in order to determine if the uncertainty in the absolute rates needed further consideration and testing in our model. The Absolute rates tested were between ranges that have been biologically recorded and observed through SPR analysis. As previously shown in Figure 2A, S1, and S3, changing the absolute rates did not significantly affect the results of the system, so rates close to the magnitude of previously observed SPR experiments were selected. The dissociation constant, on-rate, and off-rate for reaction 2 were found in Radaev et al., 2010 publication.

*Remaining reaction justifications for NRR model of TGF-β3*

Reaction biophysical data available and selected for the No Receptor Recruitment model of TGF-β3 can be found in Supplemental Table 3. The dissociation constant, on and off-rates for reaction 1 were selected from Huang 2011 publication based off of expertise knowledge as well as the latest and most abundant biophysical data available. The dissociation constant for reaction 2 is 2400 nM taken from Radaev et al., 2010. Since the absolute rates (on and off-rates) were not published with the dissociation constant, a screen was run in order to determine if the uncertainty in the absolute rates needed further consideration and testing in our model. The values for the rates between the three ligand systems changed, but the fold decrease in the absolute rates were held constant for ease in RMSE analysis. As previously shown in Figure 2A, S1, and S3, changing the absolute rates did not significantly affect the results of the system, so rates close to the magnitude of previously observed SPR experiments were selected.

Supplemental Table 1: Available SPR data for No Receptor Recruitment model of TGF-β1 ligand. The “A”, “B”, and “C” labels in the chart point to the specific source or test the data was obtained from per row. Bolded rates in reaction 1 are the values used in the final model.

| **No Receptor Recruitment Model: TGF-β1** | | | | | |
| --- | --- | --- | --- | --- | --- |
| # | Reaction | On-rate (nM^-1^s^-1^) | Off-rate (**s^-1^)** | Source | K_D_ (nM) |
| 1 | TGF-β1 + TβRII ↔  TGF-β1/TβRII | A) 1.16x10^-3^  **C)7.2x10^-4^** | A) 0.22  **C) 0.121** | A) Radaev 2010: Table 2  B) Groppe 2008: Table S1  C)Huang 2014: Table3 | A) 190  B) 390  **C)170** |
| 2 | TGF-β1 + TβRI ↔ TGF-β1/TβRI | A) 1.73x10^-6^ | A) 0.121 | A) Screened  B) Radaev 2010: Table2 | B) 70000 |
| 3 | TGF-β1/TβRII + TβRII ↔  TGF-β1/TβRII/TβRII |  |  | Homologous to Rxn1 |  |
| 4 | TGF-β1/TβRII + TβRI ↔  TGF-β1/TβRII/TβRI |  |  | Homologous to Rxn 2 |  |
| 5 | TGF-β1/TβRI + TβRII ↔ TGF-β1/TβRII/TβRI |  |  | Homologous to Rxn1 |  |
| 6 | TGF-β1/TβRI + TβRI ↔ TGF-β1/TβRI/TβRI |  |  | Homologous to Rxn2 |  |
| 7 | TGF-β1/TβRII/TβRII + TβRI ↔  TGF-β1/TβRII/TβRII/TβRI |  |  | Homologous to Rxn2 |  |
| 8 | TGF-β1/TβRII/TβRI + TβRII ↔  TGF-β1/TβRII/TβRII/TβRI |  |  | Homologous to Rxn1 |  |
| 9 | TGF-β1/TβRII/TβRI + TβRI ↔  TGF-β1/TβRII/TβRI/TβRI |  |  | Homologous to Rxn2 |  |
| 10 | TGF-β1/TβRI/TβRI + TβRII ↔ TGF-β1/TβRII/TβRI/TβRI |  |  | Homologous to Rxn1 |  |
| 11 | TGF-β1/TβRII/TβRII/TβRI + TβRI ↔ TGF-β1/TβRII/TβRII/TβRI/TβRI |  |  | Homologous to Rxn2 |  |
| 12 | TGF-β1/TβRII/TβRI/TβRI + TβRII ↔ TGF-β1/TβRII/TβRII/TβRI/TβRI |  |  | Homologous to Rxn1 |  |
| 13 | BG+TGF-β1 ↔ TGF-β1/BG | A) 1.5 x 10^-3^ | A) 7.6 x10^-4^ | A) Hinck 2018: Table 1 | A) 0.51 |
| 14 | TGF-β1/BG + TβRII ↔  TGF-β1/BG/TβRII | A) 2.24 x 10^-4^ | B) 0.24 | A) Calculated in paper  B) Radaev 2010: Table S2  C) Villarreal 2016: Table 2 | C)1070 |
| 15 | TGF-β1/TβRII + BG ↔ TGF-β1/BG/TβRII |  |  | Homologous to Rxn13 |  |
| 16 | TGF-β1/BG/TβRII + TβRI ↔  TGF-β1/BG/TβRII/TβRI |  |  | Homologous to Rxn2 |  |
| 17 | TGF-β1/TβRII/TβRI + BG ↔ TGF-β1/BG/TβRII/TβRI | A) 3.3 x 10^-5^ | A) 2.9 x10^-3^ | A) Kim 2019: Table1 | 90 |

Supplemental Table 2: Available SPR data for No Receptor Recruitment model of TGF-β2 ligand. The “A”, “B”, and “C” labels in the chart point to the specific source the data was obtained from per row. The bolded rates in reaction 1 are the values used in the final model.

| **No Receptor Recruitment Model: TGF-β2** | | | | | |
| --- | --- | --- | --- | --- | --- |
| # | Reaction | On-rate (nM^-1^s^-1^) | Off-rate (**s^-1^)** | Source | K_D_ (nM) |
| 1 | TGF-β2 + TβRII ↔  TGF-β2/TβRII | C) 4.9 x 10^-5^  **D) 4.9 x 10^-5^** | C) 1.10  **D) 0.2554** | A) Villarreal 2016: Table 5  B) Groppe 2008: Table S1  C) Radaev 2010: Table 2  D) Screened | **A) 4600**  B) 23000  C) 22449 |
| 2 | TGF-β2 + TβRI ↔ TGF-β2/TβRI | A) 9.6 x 10^-5^ | A) 1.08 | A) Radaev 2010: Table 2 | A) 11250 |
| 3 | TGF-β2/TβRII + TβRII ↔  TGF-β2/TβRII/TβRII |  |  | Homologous to Rxn1 |  |
| 4 | TGF-β2/TβRII + TβRI ↔  TGF-β2/TβRII/TβRI |  |  | Homologous to Rxn 2 |  |
| 5 | TGF-β2/TβRI + TβRII ↔ TGF-β2/TβRII/TβRI |  |  | Homologous to Rxn1 |  |
| 6 | TGF-β2/TβRI + TβRI ↔ TGF-β2/TβRI/TβRI |  |  | Homologous to Rxn2 |  |
| 7 | TGF-β2/TβRII/TβRII + TβRI ↔  TGF-β2/TβRII/TβRII/TβRI |  |  | Homologous to Rxn2 |  |
| 8 | TGF-β2/TβRII/TβRI + TβRII ↔  TGF-β2/TβRII/TβRII/TβRI |  |  | Homologous to Rxn1 |  |
| 9 | TGF-β2/TβRII/TβRI + TβRI ↔  TGF-β2/TβRII/TβRI/TβRI |  |  | Homologous to Rxn2 |  |
| 10 | TGF-β2/TβRI/TβRI + TβRII ↔ TGF-β2/TβRII/TβRI/TβRI |  |  | Homologous to Rxn1 |  |
| 11 | TGF-β2/TβRII/TβRII/TβRI + TβRI ↔ TGF-β2/TβRII/TβRII/TβRI/TβRI |  |  | Homologous to Rxn2 |  |
| 12 | TGF-β2/TβRII/TβRI/TβRI + TβRII ↔ TGF-β2/TβRII/TβRII/TβRI/TβRI |  |  | Homologous to Rxn1 |  |
| 13 | BG+TGF-β2 ↔ TGF-β2/BG | A) 1.5 x 10^-3^ | A) 7.6 x10^-4^ | A) Hinck 2018: Table 1 | A) 0.51 |
| 14 | TGF-β2/BG + TβRII ↔  TGF-β2/BG/TβRII | A) 2.24 x 10^-4^ | B) 0.24 | A) Calculated in this paper  B) Radaev 2010: Table S2  C) Villarreal 2016: Table 2 | C)1070 |
| 15 | TGF-β2/TβRII + BG ↔ TGF-β2/BG/TβRII |  |  | Homologous to Rxn13 |  |
| 16 | TGF-β2/BG/TβRII + TβRI ↔  TGF-β2/BG/TβRII/TβRI |  |  | Homologous to Rxn2 |  |
| 17 | TGF-β2/TβRII/TβRI + BG ↔ TGF-β2/BG/TβRII/TβRI | A) 3.3 x 10^-5^ | A) 2.9 x10^-3^ | A) Kim 2019: Table1 | 90 |

Supplemental Table 3: Available SPR data for No Receptor Recruitment model of TGF-β3 ligand. The “A”, “B”, and “C” labels in the chart point to the specific source the data was obtained from per row. Bolded rates in reaction 1 are the values used in the final model.

| **No Receptor Recruitment Model: TGF-β3** | | | | | |
| --- | --- | --- | --- | --- | --- |
| # | Reaction | On-rate (nM^-1^s^-1^) | Off-rate (**s^-1^)** | Source | K_D_ (nM) |
| 1 | TGF-β3 + TβRII ↔  TGF-β3/TβRII | **A) 7.4x10^-4^**  C) 1.8x10^-3^ | **A) 0.10**  C) 0.24 | A) Huang 2011: Table1  B) Groppe 2008: TableS1  C) Radaev 2010: Table2 | **A) 140**  B) 520  C) 140 |
| 2 | TGF-β3 + TβRI ↔ TGF-β3/TβRI | A) 4.167x10^-5^ | A) 0.10 | A) Screened  B) Radaev 2010: Table2 | B) 2400 |
| 3 | TGF-β3/TβRII + TβRII ↔  TGF-β3/TβRII/TβRII |  |  | Homologous to Rxn1 |  |
| 4 | TGF-β3/TβRII + TβRI ↔  TGF-β3/TβRII/TβRI |  |  | Homologues to Rxn 2 |  |
| 5 | TGF-β3/TβRI + TβRII ↔ TGF-β3/TβRII/TβRI |  |  | Homologous to Rxn1 |  |
| 6 | TGF-β3/TβRI + TβRI ↔ TGF-β3/TβRI/TβRI |  |  | Homologous to Rxn2 |  |
| 7 | TGF-β3/TβRII/TβRII + TβRI ↔  TGF-β3/TβRII/TβRII/TβRI |  |  | Homologous to Rxn2 |  |
| 8 | TGF-β3/TβRII/TβRI + TβRII ↔  TGF-β3/TβRII/TβRII/TβRI |  |  | Homologous to Rxn1 |  |
| 9 | TGF-β3/TβRII/TβRI + TβRI ↔  TGF-β3/TβRII/TβRI/TβRI |  |  | Homologous to Rxn2 |  |
| 10 | TGF-β3/TβRI/TβRI + TβRII ↔ TGF-β3/TβRII/TβRI/TβRI |  |  | Homologous to Rxn1 |  |
| 11 | TGF-β3/TβRII/TβRII/TβRI + TβRI ↔ TGF-β3/TβRII/TβRII/TβRI/TβRI |  |  | Homologous to Rxn2 |  |
| 12 | TGF-β3/TβRII/TβRI/TβRI + TβRII ↔ TGF-β3/TβRII/TβRII/TβRI/TβRI |  |  | Homologous to Rxn1 |  |
| 13 | BG+TGF-β3 ↔ TGF-β3/BG | A) 1.5 x 10^-3^ | A) 7.6 x10^-4^ | A) Hinck 2018: Table 1 | A) 0.51 |
| 14 | TGF-β3/BG + TβRII ↔  TGF-β3/BG/TβRII | A) 2.24 x 10^-4^ | B) 0.24 | A) Calculated in this paper  B) Radaev 2010: Table S2  C) Villarreal 2016: Table 2 | C)1070 |
| 15 | TGF-β3/TβRII + BG ↔ TGF-β3/BG/TβRII |  |  | Homologous to Rxn13 |  |
| 16 | TGF-β3/BG/TβRII + TβRI ↔  TGF-β3/BG/TβRII/TβRI |  |  | Homologous to Rxn2 |  |
| 17 | TGF-β3/TβRII/TβRI + BG ↔ TGF-β3/BG/TβRII/TβRI | A) 3.3 x 10^-5^ | A) 2.9 x10^-3^ | A) Kim 2019: Table1 | A) 90 |

*S6. Single-stage Receptor Recruitment model creation*

The justifications for the homologous reactions, screens, and reaction kinetics in the Single-stage Receptor Recruitment (SRR) model are the same as the NRR model except for the added receptor recruitment of TβRI by a ligand bound TβRII found in literature that affects reactions 4, 7,11, and 16 (Supplemental Table 4-6). Reactions 7,11, and 16 are homologous to reaction 4 because there is an addition of a single TβRI when a ligand bound TβRII is present and not already bound to TβRI. The justifications and kinetic rates for reactions 13 through 17 in the SRR model are still the same as the NRR model, but reaction 16’s kinetics changed due to the homology to reaction 4 as mentioned previously.

*Remaining reaction justifications for SRR model of TGF-β1*

Reaction biophysical data available and selected for the SRR model of TGF-β1 can be found in Supplemental Table 4. The same absolute rate screen for reaction 2 carried out in the NRR model was performed in the SRR model. As previously shown in Figure 2A, S1, and S3, changing the absolute rates did not significantly affect the results of the system, so the same fold decrease in absolute rates chosen in the NRR model were also chosen for the SRR model to maintain comparison integrity. The dissociation constant, on and off-rates for reaction 4 were selected from Huang et al., 2014 publication based off of expertise knowledge as well as the latest and most abundant biophysical data available (Supplemental Table 4).

*Remaining reaction justifications for SRR model of TGF-β2*

Reaction biophysical data available and selected for the SRR model of TGF-β2 can be found in Supplemental Table 5. The same absolute rate screen for reaction 1 carried out in the NRR model was performed in the SRR model. As previously shown in Figure 2A, S1, and S3, changing the absolute rates did not significantly affect the results of the system, so the same rates chosen in the NRR model were also chosen for the SRR model to maintain comparison integrity. The dissociation constant, on-rate, and off-rate for reaction 4 were selected from Radaev et al., 2010 publication based off of expertise knowledge as well as the latest and most abundant biophysical data available in a single publication (Supplemental Table 5).

*Remaining reaction justifications for SRR model of TGF-β3*

Reaction biophysical data available and selected for the SRR model of TGF-β3 can be found in Supplemental Table 6. The same absolute rate screen for reaction 2 carried out in the NRR model was performed in the SRR model. As previously shown in Figure 2A, S1, and S3, changing the absolute rates did not significantly affect the results of the system, so the same rates chosen in the NRR model were also chosen for the SRR model to maintain comparison integrity. The dissociation constant, on and off-rates for reaction 4 were selected from Huang et al., 2011 publication based off of expertise knowledge as well as the latest and most abundant biophysical data available (Supplemental Table 6).

Supplemental Table 4: Available SPR data for Single-stage Receptor Recruitment model of TGF-β1 ligand. The “A”, “B”, and “C” labels in the chart point to the specific source or test the data was obtained from per row. The rows highlighted in red show the reactions changed when adding the TβRII Recruitment of TβRI of TβRI by TβRII. Bolded rates in reaction 4 are the values used in the final model.

| **Single-stage Receptor Recruitment Model: TGF-β1** | | | | | |
| --- | --- | --- | --- | --- | --- |
| # | Reaction | On-rate (nM^-1^s^-1^) | Off-rate (**s^-1^)** | Source | K_D_ (nM) |
| 1 | TGF-β1 + TβRII ↔  TGF-β1/TβRII | A) 7.2x10^-4^ | A) 0.121 | A) Huang 2014: Table3 | A) 170 |
| 2 | TGF-β1 + TβRI ↔ TGF-β1/TβRI | A) 1.73x10^-6^ | A) 0.121 | A) Screened  B) Radaev 2010: Table2 | B) 70000 |
| 3 | TGF-β1/TβRII + TβRII ↔  TGF-β1/TβRII/TβRII |  |  | Homologous to Rxn1 |  |
| 4 | TGF-β1/TβRII + TβRI ↔  TGF-β1/TβRII/TβRI | A) 9.7x10^-5^  **C)3.3x10^-5^** | A) 6.8x10^-3^  **C)7.6x10^-3^** | A) Radaev 2010: Table2  B) Groppe 2008: Table S1  C)Huang 2014:Table3 | A) 70  B) 2530  **C) 240** |
| 5 | TGF-β1/TβRI + TβRII ↔ TGF-β1/TβRII/TβRI |  |  | Homologous to Rxn1 |  |
| 6 | TGF-β1/TβRI + TβRI ↔ TGF-β1/TβRI/TβRI |  |  | Homologous to Rxn2 |  |
| 7 | TGF-β1/TβRII/TβRII + TβRI ↔  TGF-β1/TβRII/TβRII/TβRI |  |  | Homologous to Rxn4 |  |
| 8 | TGF-β1/TβRII/TβRI + TβRII ↔  TGF-β1/TβRII/TβRII/TβRI |  |  | Homologous to Rxn1 |  |
| 9 | TGF-β1/TβRII/TβRI + TβRI ↔  TGF-β1/TβRII/TβRI/TβRI |  |  | Homologous to Rxn2 |  |
| 10 | TGF-β1/TβRI/TβRI + TβRII ↔ TGF-β1/TβRII/TβRI/TβRI |  |  | Homologous to Rxn1 |  |
| 11 | TGF-β1/TβRII/TβRII/TβRI + TβRI ↔ TGF-β1/TβRII/TβRII/TβRI/TβRI |  |  | Homologous to Rxn4 |  |
| 12 | TGF-β1/TβRII/TβRI/TβRI + TβRII ↔ TGF-β1/TβRII/TβRII/TβRI/TβRI |  |  | Homologous to Rxn1 |  |
| 13 | BG+TGF-β1 ↔ TGF-β1/BG | A) 1.5 x 10^-3^ | A) 7.6 x10^-4^ | A) Hinck 2018: Table 1 | A) 0.51 |
| 14 | TGF-β1/BG + TβRII ↔  TGF-β1/BG/TβRII | A) 2.24 x 10^-4^ | B) 0.24 | A) Calculated in paper  B) Radaev 2010: Table S2  C) Villarreal 2016: Table 2 | C)1070 |
| 15 | TGF-β1/TβRII + BG ↔ TGF-β1/BG/TβRII |  |  | Homologous to Rxn13 |  |
| 16 | TGF-β1/BG/TβRII + TβRI ↔  TGF-β1/BG/TβRII/TβRI |  |  | Homologous to Rxn4 |  |
| 17 | TGF-β1/TβRII/TβRI + BG ↔ TGF-β1/BG/TβRII/TβRI | A) 3.3 x 10^-5^ | A) 2.9 x10^-3^ | A) Kim 2019: Table1 | 90 |

Supplemental Table 5: Available SPR data for Single-stage Receptor Recruitment model of TGF-β2 ligand. The “A”, “B”, and “C” labels in the chart point to the specific source the data was obtained from per row. The rows highlighted in red show the reactions changed when adding the experimentally determined recruitment of TβRI by TβRII. The bolded rates in reaction 4 are the values used in the final model.

| **Single-stage Receptor Recruitment Model: TGF-β2** | | | | | |
| --- | --- | --- | --- | --- | --- |
| # | Reaction | On-rate (nM^-1^s^-1^) | Off-rate (**s^-1^)** | Source | K_D_ (nM) |
| 1 | TGF-β2 + TβRII ↔  TGF-β2/TβRII | A) 4.9 x 10^-5^ | A) 0.2554 | A) Screened  B) Villarreal 2016: Table 5 | B) 4600 |
| 2 | TGF-β2 + TβRI ↔ TGF-β2/TβRI | A) 9.6 x 10^-5^ | A) 1.08 | A) Radaev 2010: Table 2 | A) 11250 |
| 3 | TGF-β2/TβRII + TβRII ↔  TGF-β2/TβRII/TβRII |  |  | Homologous to Rxn1 |  |
| 4 | TGF-β2/TβRII + TβRI ↔  TGF-β2/TβRII/TβRI | **A) 1.8 x 10^-4^** | **A) 2.9 x 10^-3^** | A)Radaev 2010: Table 2  B)Groppe 2008:Table S1 | **A) 16**  B) 1170 |
| 5 | TGF-β2/TβRI + TβRII ↔ TGF-β2/TβRII/TβRI |  |  | Homologous to Rxn1 |  |
| 6 | TGF-β2/TβRI + TβRI ↔ TGF-β2/TβRI/TβRI |  |  | Homologous to Rxn2 |  |
| 7 | TGF-β2/TβRII/TβRII + TβRI ↔  TGF-β2/TβRII/TβRII/TβRI |  |  | Homologous to Rxn4 |  |
| 8 | TGF-β2/TβRII/TβRI + TβRII ↔  TGF-β2/TβRII/TβRII/TβRI |  |  | Homologous to Rxn1 |  |
| 9 | TGF-β2/TβRII/TβRI + TβRI ↔  TGF-β2/TβRII/TβRI/TβRI |  |  | Homologous to Rxn2 |  |
| 10 | TGF-β2/TβRI/TβRI + TβRII ↔ TGF-β2/TβRII/TβRI/TβRI |  |  | Homologous to Rxn1 |  |
| 11 | TGF-β2/TβRII/TβRII/TβRI + TβRI ↔ TGF-β2/TβRII/TβRII/TβRI/TβRI |  |  | Homologous to Rxn4 |  |
| 12 | TGF-β2/TβRII/TβRI/TβRI + TβRII ↔ TGF-β2/TβRII/TβRII/TβRI/TβRI |  |  | Homologous to Rxn1 |  |
| 13 | BG+TGF-β2 ↔ TGF-β2/BG | A) 1.5 x 10^-3^ | A) 7.6 x10^-4^ | A) Hinck 2018: Table 1 | A) 0.51 |
| 14 | TGF-β2/BG + TβRII ↔  TGF-β2/BG/TβRII | A) 2.24 x 10^-4^ | B) 0.24 | A) Calculated in this paper  B) Radaev 2010: Table S2  C) Villarreal 2016: Table 2 | C)1070 |
| 15 | TGF-β2/TβRII + BG ↔ TGF-β2/BG/TβRII |  |  | Homologous to Rxn13 |  |
| 16 | TGF-β2/BG/TβRII + TβRI ↔  TGF-β2/BG/TβRII/TβRI |  |  | Homologous to Rxn4 |  |
| 17 | TGF-β2/TβRII/TβRI + BG ↔ TGF-β2/BG/TβRII/TβRI | A) 3.3 x 10^-5^ | A) 2.9 x10^-3^ | A) Kim 2019: Table1 | 90 |

Supplemental Table 6: Available SPR data for Single-stage Receptor Recruitment model of TGF-β3 ligand. The “A”, “B”, and “C” labels in the chart point to the specific source the data was obtained from per row. The rows highlighted in red show the reactions changed when adding the experimentally determined recruitment of TβRI by TβRII. The bolded rates in reaction 4 are the values used in the final model.

| **Single-stage Receptor Recruitment Model: TGF-β3** | | | | | |
| --- | --- | --- | --- | --- | --- |
| # | Reaction | On-rate (nM^-1^s^-1^) | Off-rate (**s^-1^)** | Source | K_D_ (nM) |
| 1 | TGF-β3 + TβRII ↔  TGF-β3/TβRII | A) 7.4x10^-4^ | A) 0.10 | A) Huang 2011: Table1 | A) 140 |
| 2 | TGF-β3 + TβRI ↔ TGF-β3/TβRI | A) 4.167x10^-5^ | A) 0.10 | A) Screened  B) Radaev 2010: Table2 | B) 2400 |
| 3 | TGF-β3/TβRII + TβRII ↔  TGF-β3/TβRII/TβRII |  |  | Homologous to Rxn1 |  |
| 4 | TGF-β3/TβRII + TβRI ↔  TGF-β3/TβRII/TβRI | **A) 3.5x10^-5^**  C) 9.6x10^-5^ | **A) 1.2x10^-3^**  C) 1.3x10^-3^ | A) Huang 2011: Table 1  B) Groppe 2008: Table S1  C) Radaev 2010: Table2 | **A) 34**  B) 600  C) 14.0 |
| 5 | TGF-β3/TβRI + TβRII ↔ TGF-β3/TβRII/TβRI |  |  | Homologous to Rxn1 |  |
| 6 | TGF-β3/TβRI + TβRI ↔ TGF-β3/TβRI/TβRI |  |  | Homologous to Rxn2 |  |
| 7 | TGF-β3/TβRII/TβRII + TβRI ↔  TGF-β3/TβRII/TβRII/TβRI |  |  | Homologous to Rxn4 |  |
| 8 | TGF-β3/TβRII/TβRI + TβRII ↔  TGF-β3/TβRII/TβRII/TβRI |  |  | Homologous to Rxn1 |  |
| 9 | TGF-β3/TβRII/TβRI + TβRI ↔  TGF-β3/TβRII/TβRI/TβRI |  |  | Homologous to Rxn2 |  |
| 10 | TGF-β3/TβRI/TβRI + TβRII ↔ TGF-β3/TβRII/TβRI/TβRI |  |  | Homologous to Rxn1 |  |
| 11 | TGF-β3/TβRII/TβRII/TβRI + TβRI ↔ TGF-β3/TβRII/TβRII/TβRI/TβRI |  |  | Homologous to Rxn4 |  |
| 12 | TGF-β3/TβRII/TβRI/TβRI + TβRII ↔ TGF-β3/TβRII/TβRII/TβRI/TβRI |  |  | Homologous to Rxn1 |  |
| 13 | BG+TGF-β3 ↔ TGF-β3/BG | A) 1.5 x 10^-3^ | A) 7.6 x10^-4^ | A) Hinck 2018: Table 1 | A) 0.51 |
| 14 | TGF-β3/BG + TβRII ↔  TGF-β3/BG/TβRII | A) 2.24 x 10^-4^ | B) 0.24 | A) Calculated in this paper  B) Radaev 2010: Table S2  C) Villarreal 2016: Table 2 | C)1070 |
| 15 | TGF-β3/TβRII + BG ↔ TGF-β3/BG/TβRII |  |  | Homologous to Rxn13 |  |
| 16 | TGF-β3/BG/TβRII + TβRI ↔  TGF-β3/BG/TβRII/TβRI |  |  | Homologous to Rxn4 |  |
| 17 | TGF-β3/TβRII/TβRI + BG ↔ TGF-β3/BG/TβRII/TβRI | A) 3.3 x 10^-5^ | A) 2.9 x10^-3^ | A) Kim 2019: Table1 | 90 |

*S7. Two-stage Receptor Recruitment model creation*

The Two-stage Receptor Recruitment (TRR) model continues to build off of the NRR and SRR models. The justifications for the homologous reactions and reaction kinetics are the same as the SRR model except for the added receptor recruitment of TβRII by a ligand bound TβRI that affects reactions 5, 10, and 12 (Supplemental Table 7-9). Reactions 10 and 12 are homologous to reaction 5 because there is an addition of a single TβRII when a ligand bound TβRI is present and not previously bound to a TβRII.

*Remaining reaction justifications for TRR model for TGF-β1, TGF-β2, and TGF-β3*

Reaction biophysical data available and selected for the TRR model for TGF-β1, TGF-β2, TGF-β3 can be found in Supplemental Tables 7, 8, and 9, respectfully. Screens were run to determine the optimal degree of recruitment for TβRI on TβRII (range of 1 to 700 degrees of recruitment) and to determine the effect of changing the absolute rates (range of 1 to 500-fold decrease in absolute rates) on all three ligand systems. The values for the rates between the three ligand systems changed, but the fold decrease in the absolute rates were held constant for the RMSE analysis, these value differences can be found in the supplemental tables. As previously shown in Figures 2B and 3S, simultaneously decreasing the magnitude of the absolute rates did not appreciably affect the RMSE analysis. Due to the minimal impact of changing the absolute rates simultaneously, the absolute rates selected were close to the magnitude of previously observed SPR experiments. Next, a more detailed screen on the degree of receptor recruitment of reactions 5, 10, and 12 were tested as shown in Figure 4S. A degree recruitment of roughly 5 across all of the ligand systems produces the lowest RMSE, therefore, the dissociation constant chosen was five-fold more favorable than reaction 1’s dissociation constant in each of the ligand systems (Supplemental Table 7, 8, and 9).

Supplemental Table 7: Available SPR data for Two-stage Receptor Recruitment model of TGF-β1 ligand. The “A”, “B”, and “C” labels in the chart point to the specific source or test the data was obtained from per row. The rows highlighted in red show the reactions changed when adding the experimentally determined recruitment of TβRI by TβRII. The rows highlighted in blue show the reactions affected when adding the theorized recruitment of TβRII by TβRI.

| **Two-stage Receptor Recruitment Model: TGF-β1** | | | | | |
| --- | --- | --- | --- | --- | --- |
| # | Reaction | On-rate (nM^-1^s^-1^) | Off-rate (**s^-1^)** | Source | K_D_ (nM) |
| 1 | TGF-β1 + TβRII ↔  TGF-β1/TβRII | A) 7.2x10^-4^ | A) 0.121 | A) Huang 2014: Table3 | A) 170 |
| 2 | TGF-β1 + TβRI ↔ TGF-β1/TβRI | B) 1.73x10^-6^ | B) 0.121 | A) Radaev 2010: Table2  B) Screened | A) 70000 |
| 3 | TGF-β1/TβRII + TβRII ↔  TGF-β1/TβRII/TβRII |  |  | Homologous to Rxn1 |  |
| 4 | TGF-β1/TβRII + TβRI ↔  TGF-β1/TβRII/TβRI | A) 3.3x10^-5^ | A) 7.6x10^-3^ | A) Huang 2014: Table3 | A) 240 |
| 5 | TGF-β1/TβRI + TβRII ↔ TGF-β1/TβRII/TβRI | A) 3.83547x10^-3^ | A) 0.12516 | A) Screened | A) 33 |
| 6 | TGF-β1/TβRI + TβRI ↔ TGF-β1/TβRI/TβRI |  |  | Homologous to Rxn2 |  |
| 7 | TGF-β1/TβRII/TβRII + TβRI ↔  TGF-β1/TβRII/TβRII/TβRI |  |  | Homologous to Rxn4 |  |
| 8 | TGF-β1/TβRII/TβRI + TβRII ↔TGF-β1/TβRII/TβRII/TβRI |  |  | Homologous to Rxn1 |  |
| 9 | TGF-β1/TβRII/TβRI + TβRI ↔  TGF-β1/TβRII/TβRI/TβRI |  |  | Homologous to Rxn2 |  |
| 10 | TGF-β1/TβRI/TβRI + TβRII ↔ TGF-β1/TβRII/TβRI/TβRI |  |  | Homologous to Rxn5 |  |
| 11 | TGF-β1/TβRII/TβRII/TβRI + TβRI ↔ TGF-β1/TβRII/TβRII/TβRI/TβRI |  |  | Homologous to Rxn4 |  |
| 12 | TGF-β1/TβRII/TβRI/TβRI + TβRII ↔ TGF-β1/TβRII/TβRII/TβRI/TβRI |  |  | Homologous to Rxn5 |  |
| 13 | BG+TGF-β1 ↔ TGF-β1/BG | A) 1.5 x 10^-3^ | A) 7.6 x10^-4^ | A) Hinck 2018: Table 1 | A) 0.51 |
| 14 | TGF-β1/BG + TβRII ↔  TGF-β1/BG/TβRII | A) 2.24 x 10^-4^ | B) 0.24 | A) Calculated in paper  B) Radaev 2010: Table S2  C) Villarreal 2016: Table 2 | C)1070 |
| 15 | TGF-β1/TβRII + BG ↔ TGF-β1/BG/TβRII |  |  | Homologous to Rxn13 |  |
| 16 | TGF-β1/BG/TβRII + TβRI ↔  TGF-β1/BG/TβRII/TβRI |  |  | Homologous to Rxn4 |  |
| 17 | TGF-β1/TβRII/TβRI + BG ↔TGF-β1/BG/TβRII/TβRI | A) 3.3 x 10^-5^ | A) 2.9 x10^-3^ | A) Kim 2019: Table1 | 90 |

Supplemental Table 8: Available SPR data for Two-stage Receptor Recruitment model of TGF-β2 ligand. The “A”, “B”, and “C” labels in the chart point to the specific source the data was obtained from per row. The rows highlighted in red show the reactions changed when adding the experimentally determined recruitment of TβRI by TβRII. The rows highlighted in blue show the reactions affected when adding the theorized recruitment of TβRII by TβRI.

| **Two-stage Receptor Recruitment Model: TGF-β2** | | | | | |
| --- | --- | --- | --- | --- | --- |
| # | Reaction | On-rate (nM^-1^s^-1^) | Off-rate (**s^-1^)** | Source | K_D_ (nM) |
| 1 | TGF-β2 + TβRII ↔  TGF-β2/TβRII | A) 4.9 x 10^-5^ | A) 0.2554 | A) Screened  B) Villarreal 2016:Table 5 | B) 4600 |
| 2 | TGF-β2 + TβRI ↔ TGF-β2/TβRI | A) 9.6 x 10^-5^ | A) 1.08 | A) Radaev 2010: Table 2 | A) 11250 |
| 3 | TGF-β2/TβRII + TβRII ↔  TGF-β2/TβRII/TβRII |  |  | Homologous to Rxn1 |  |
| 4 | TGF-β2/TβRII + TβRI ↔  TGF-β2/TβRII/TβRI | A) 1.8 x 10^-4^ | A) 2.9 x 10^-3^ | A) Radaev 2010: Table 2 | A) 16 |
| 5 | TGF-β2/TβRI + TβRII ↔ TGF-β2/TβRII/TβRI | A) 4.9 x 10^-5^ | A) 0.0438 | A) Screened | A) 893 |
| 6 | TGF-β2/TβRI + TβRI ↔ TGF-β2/TβRI/TβRI |  |  | Homologous to Rxn2 |  |
| 7 | TGF-β2/TβRII/TβRII + TβRI ↔  TGF-β2/TβRII/TβRII/TβRI |  |  | Homologous to Rxn4 |  |
| 8 | TGF-β2/TβRII/TβRI + TβRII ↔  TGF-β2/TβRII/TβRII/TβRI |  |  | Homologous to Rxn1 |  |
| 9 | TGF-β2/TβRII/TβRI + TβRI ↔  TGF-β2/TβRII/TβRI/TβRI |  |  | Homologous to Rxn2 |  |
| 10 | TGF-β2/TβRI/TβRI + TβRII ↔ TGF-β2/TβRII/TβRI/TβRI |  |  | Homologous to Rxn5 |  |
| 11 | TGF-β2/TβRII/TβRII/TβRI + TβRI ↔ TGF-β2/TβRII/TβRII/TβRI/TβRI |  |  | Homologous to Rxn4 |  |
| 12 | TGF-β2/TβRII/TβRI/TβRI + TβRII ↔ TGF-β2/TβRII/TβRII/TβRI/TβRI |  |  | Homologous to Rxn5 |  |
| 13 | BG+TGF-β2 ↔ TGF-β2/BG | A) 1.5 x 10^-3^ | A) 7.6 x10^-4^ | A) Hinck 2018: Table 1 | A) 0.51 |
| 14 | TGF-β2/BG + TβRII ↔  TGF-β2/BG/TβRII | A) 2.24 x 10^-4^ | B) 0.24 | A) Calculated in this paper  B) Radaev 2010: Table S2  C) Villarreal 2016: Table 2 | C)1070 |
| 15 | TGF-β2/TβRII + BG ↔ TGF-β2/BG/TβRII |  |  | Homologous to Rxn13 |  |
| 16 | TGF-β2/BG/TβRII + TβRI ↔  TGF-β2/BG/TβRII/TβRI |  |  | Homologous to Rxn4 |  |
| 17 | TGF-β2/TβRII/TβRI + BG ↔ TGF-β2/BG/TβRII/TβRI | A) 3.3 x 10^-5^ | A) 2.9 x10^-3^ | A) Kim 2019: Table1 | 90 |

Supplemental Table 9: Available SPR data for Two-stage Receptor Recruitment model of TGF-β3 ligand. The “A”, “B”, and “C” labels in the chart point to the specific source the data was obtained from per row. The rows highlighted in red show the reactions changed when adding the experimentally determined recruitment of TβRI by TβRII. The rows highlighted in blue show the reactions affected when adding the theorized recruitment of TβRII by TβRI.

| **Two-stage Receptor Recruitment Model: TGF-β3** | | | | | |
| --- | --- | --- | --- | --- | --- |
| # | Reaction | On-rate (nM^-1^s^-1^) | Off-rate (**s^-1^)** | Source | K_D_ (nM) |
| 1 | TGF-β3 + TβRII ↔  TGF-β3/TβRII | A) 7.4x10^-4^ | A) 0.10 | A) Huang 2011: Table1 | A) 140 |
| 2 | TGF-β3 + TβRI ↔ TGF-β3/TβRI | A) 4.167x10^-5^ | A) 0.10 | A) Screened  B) Radaev 2010: Table2 | B) 2400 |
| 3 | TGF-β3/TβRII + TβRII ↔  TGF-β3/TβRII/TβRII |  |  | Homologous to Rxn1 |  |
| 4 | TGF-β3/TβRII + TβRI ↔  TGF-β3/TβRII/TβRI | A) 3.5x10^-5^ | A) 1.2x10^-3^ | A) Huang 2011: Table 1 | A) 34 |
| 5 | TGF-β3/TβRI + TβRII ↔ TGF-β3/TβRII/TβRI | A) 3.94x10^-3^ | A) 0.103 | A) Screened | A) 27 |
| 6 | TGF-β3/TβRI + TβRI ↔ TGF-β3/TβRI/TβRI |  |  | Homologous to Rxn2 |  |
| 7 | TGF-β3/TβRII/TβRII + TβRI ↔  TGF-β3/TβRII/TβRII/TβRI |  |  | Homologous to Rxn4 |  |
| 8 | TGF-β3/TβRII/TβRI + TβRII ↔  TGF-β3/TβRII/TβRII/TβRI |  |  | Homologous to Rxn1 |  |
| 9 | TGF-β3/TβRII/TβRI + TβRI ↔  TGF-β3/TβRII/TβRI/TβRI |  |  | Homologous to Rxn2 |  |
| 10 | TGF-β3/TβRI/TβRI + TβRII ↔ TGF-β3/TβRII/TβRI/TβRI |  |  | Homologous to Rxn5 |  |
| 11 | TGF-β3/TβRII/TβRII/TβRI + TβRI ↔ TGF-β3/TβRII/TβRII/TβRI/TβRI |  |  | Homologous to Rxn4 |  |
| 12 | TGF-β3/TβRII/TβRI/TβRI + TβRII ↔ TGF-β3/TβRII/TβRII/TβRI/TβRI |  |  | Homologous to Rxn5 |  |
| 13 | BG+TGF-β3 ↔ TGF-β3/BG | A) 1.5 x 10^-3^ | A) 7.6 x10^-4^ | A) Hinck 2018: Table 1 | A) 0.51 |
| 14 | TGF-β3/BG + TβRII ↔  TGF-β3/BG/TβRII | A) 2.24 x 10^-4^ | B) 0.24 | A) Calculated in this paper  B) Radaev 2010: Table S2  C) Villarreal 2016: Table 2 | C)1070 |
| 15 | TGF-β3/TβRII + BG ↔ TGF-β3/BG/TβRII |  |  | Homologous to Rxn13 |  |
| 16 | TGF-β3/BG/TβRII + TβRI ↔  TGF-β3/BG/TβRII/TβRI |  |  | Homologous to Rxn4 |  |
| 17 | TGF-β3/TβRII/TβRI + BG ↔ TGF-β3/BG/TβRII/TβRI | A) 3.3 x 10^-5^ | A) 2.9 x10^-3^ | A) Kim 2019: Table1 | 90 |

*S8. Receptor justification*

From Wakefield et al., 1987, a median value of 10,000 is used for TGF-β receptors found per epithelial cell. Equimolar receptor concentrations were applied, 5000 Type I and Type II receptors with 10% at the surface of cell membrane (Vilar et al., 2006; Di Gulilenio et al. 2003; Chung et al., 2009). Therefore, the starting point for the simulations will be 500 Type I and II receptors which is equal to about 160 nM with the cell volume determined in Equation 3.

Equation 3:

$$Volume=10.2\mu m\times10.2\mu m\times0.05 \mu m= 5.2{\mu m}^{3}=5.2\times{{10}^{-6}}^{3}m^{3}=5.2\times{10}^{-18}m^{3}=5.2\times{10}^{-18}\times\left( \frac{1}{0.001} \right)Litre=5.2\times{10}^{-15}Litre$$

$$conversion=\frac{500 molecules}{(\left( 6.022\times{10}^{23} molecules \right)\times5.2\times{10}^{-15}Litre)}\times1\frac{Mole}{Litre}=159.67nM$$

The volume selected was a value similar to those used in other computational models of growth factors (Karim et al., 2012) and was informed by recorded data for the diameter size of common epithelial cells and the size of epithelial cells apical membrane (Devalia et al., 1990; Mitra et al., 2004). Although maintaining a volume close to the proposed extracellular space in question is important to drawing biological conclusions, narrowing in on the exact volume is time exhaustive because changing the volume does not significantly alter the signaling pattern. If the receptor concentration is held constant, decreasing the simulated extracellular volume around the cell will increase the number of receptors per milliliter (mL). This volume change will lead to a model more sensitive to detecting signaling patterns at lower receptor levels but altering the volume does not significantly change the trend of the signaling pattern. The trends of the simulation results are more dependent on the ratios of receptor to ligand levels and BG to receptor levels which are tested and analyzed in this paper.

In our computational models, endocytosis and receptor recycling were combined into one step. Drawing from the knowledge of a previous paper and expert knowledge, we used a very slow rate for this combined step (Karim et al., 2012). The TGF-β receptor complex assembly does not have a strong accumulation of higher order intermediate complexes. If the complexes were not able to dissociate when formed, this rate may need to be faster, but with the kinetic rates used for the reactions in these models, the intermediate complexes are able to dissociate freely. Including this in our model does not significantly affect the results but does make the model more biological relevant by considering endocytosis and receptor recycling.
